# Progranulin deficiency induces lipid droplet accumulation in microglia via a STAT3-GPAT3 axis

**DOI:** 10.64898/2026.01.17.700052

**Authors:** Jun Li, Yiyue Shi, Haoyuan Guan, Yifei Lu, Tuo Yi, Wei Li, Yini Wang, Yeting Guo, Bei Li, Caihong Zhu

## Abstract

Heterozygous mutations in the progranulin (PGRN) encoding gene *GRN* cause frontotemporal dementia (FTD), whereas homozygous *GRN* mutations lead to neuronal ceroid lipofuscinosis (NCL). However, the mechanisms underlying neurodegeneration due to PGRN deficiency remain unclear. In the aged brains or under neurodegenerative conditions, the accumulation of lipid droplets (LDs) in microglia contributes to cellular dysfunction and pro-inflammatory responses, exacerbating neurodegenerative pathology. Here, we investigated how PGRN deficiency induces LD accumulation in microglia. We found that PGRN ablation significantly upregulated triglycerol-3-phosphate acyltransferase 3 (GPAT3), the first and rate-limiting enzyme for triacylglycerol biosynthesis, and increased LD formation in both *Grn^-/-^* BV2 microglial cells and *Grn^-/-^*mouse brains. Mechanistic study revealed that PGRN deficiency upregulates GPAT3 via stimulating signal transduction and activator of transcription factor 3 (STAT3) signaling. Silencing *Gpat3*, inhibiting GPAT3 activity with the small molecule inhibitor FSG67, or PGRN restoration reduced LD accumulation and rescued the cytotoxicity of *Grn*^-/-^ microglia conditional medium (MCM) toward N2a cells. Notably, intraperitoneal administration of FSG67 or AAV-MG1.2-mCx3cr1-mGrn-eGFP-mediated microglia-specific gene therapy reduced microglial LDs and ameliorated behavior phenotypes in *Grn*^-/-^ mice. Our study elucidates the role of PGRN loss in microglial lipid homeostasis and identifies the STAT3-GPAT3 axis as a potential therapeutic target for LD-associated neurodegenerative diseases.

---

Progranulin (PGRN), a glycoprotein encoded by *GRN* gene, plays pleiotropic roles in the central nervous system (CNS)^1^. Genome-wide association studies (GWAS) revealed that PGRN haploinsufficiency caused by heterozygous loss-of-function (LOF) mutations in the *GRN* gene represents a major cause of frontotemporal lobar degeneration (FTLD)^2,3^, whereas homozygous *GRN* mutations leads to a rare lysosome storage disease neuronal ceroid lipofuscinosis (NCL)^4^. *GRN* polymorphisms that decrease PGRN levels have also been linked to other neurodegenerative diseases, such as Alzheimer’s disease (AD)^5,6^, Parkinson’s disease (PD)^7^ and Lewy body dementia^8^. In the CNS, PGRN is primarily expressed by microglia and neurons^9^. After translation at endoplasmic reticulum (ER) and N-glycosylation at Golgi apparatus, PGRN can be delivered to endo-lysosome or secreted to extracellular space. The secreted protein can be recycled to lysosome by directly interacting with a membrane receptor sortilin (SORT1) or indirectly associating with prosaposin (PSAP) and the mannose 6-phosphate receptor (M6PR)^10^. Within lysosome, PGRN is proteolytically cleaved into granulin peptides and plays key roles in lysosomal function^11^.

Microglia are the resident immune cells of the CNS^12^ and express the highest level of PGRN^9^. Previous studies demonstrate that PGRN is critical to maintain microglial homeostasis and suppress excessive microglial activation and neuroinflammation^13^. Upon activation during neurodegeneration or aging, microglia undergo marked morphological, transcriptional and functional changes, such as the ramified processes, altered cytokine profiles, reduced phagocytosis capacity and impaired lipid homeostasis^14^. Specifically, the lipid droplet accumulating microglia (LDAM) represent a dysfunctional and proinflammatory state during neurodegenerative and aging brains^15^.

Lipid droplets (LDs) are dynamic organelles that store hydrophobic neutral lipids, such as triacylglycerols (TAGs) and sterol esters, within a phospholipid monolayer^16^. In the CNS, LDs are crucial for lipid homeostasis, cellular metabolism, oxidative stress prevention and neuroinflammation regulation^17,18^. Although lipid accumulation in glial cells was first described by Alois Alzheimer over a century ago^19^, its role in neurodegeneration remains largely unexplored. Risk factors of AD such as APOE4, PICALM have been associated with microglial LD accumulation^20,21^. Moreover, Amyloid β (Aβ) deposits have been linked to LD-mediated microglial dysfunction^22,23^. These findings suggest that LDs represent a critical mediator for neuroinflammation and neurodegeneration. The role of PGRN in lipid homeostasis has recently been explored^24^. Lipidomic^25^ and proteomic^26^ analysis uncovered lysosomal lipid dysregulation in *Grn*^-/-^ mice. Additionally, LD accumulation has also been observed in microglia from aged as well as *Grn*^-/-^ mice^15^ and in induced microglia (iMGs) derived from FTD-*GRN* patients^27^, associated with lysosomal dysfunction, phagocytosis defects, elevated ROS and increased proinflammatory cytokines release. However, the mechanisms underlying the LD accumulation in PGRN-deficient microglia remained unknown.

Glycerol-3-phosphate acyltransferases (GPAT1-4) catalyze the first and rate-limiting step in triacylglycerol (TAG) synthesis by converting glycerol-3-phosphate and acyl-CoA to lysophosphatidic acid (LPA)^28^. GPAT1 and GPAT2 are localized on the outer mitochondrial membrane, while GPAT3 and GPAT4 are found in the endoplasmic reticulum (ER)^29^. It has been reported that triacylglycerol synthesis directed by GPAT3 and GPAT4 is required for LD formation in myeloid cells^30^. Therefore, GPAT3/4 represent a critical regulator for LD accumulation. However, whether GPATs are involved in LDAM accumulation during aging and neurodegeneration remained to be elucidated.

In this study, we demonstrate that PGRN deficiency upregulates IL-6 expression and stimulates signal transduction and activator of transcription factor 3 (STAT3) signaling, thereby increasing the expression of GPAT3 and leading to LD accumulation in *Grn^-/-^*microglia. Microglia-specific PGRN restoration normalized GPAT3 levels and reduced LDs under *Grn* knockout conditions, highlighting PGRN’s critical role in maintaining lipid homeostasis. Moreover, downregulating GPAT3 expression or inhibiting GPAT3 activity can effectively alleviate these defects, suggesting a potential therapeutic strategy for PGRN-related neurodegeneration. Our findings not only establish a PGRN-STAT3-GPAT3-LD cascade, but also provide novel insights into PGRN’s function and identify the STAT3-GPAT3 axis as a potential therapeutic target for PGRN-associated neurodegenerative diseases.

## Results

### PGRN deficiency drives lipid droplet accumulation in microglia

Prior studies have reported the involvement of PGRN in lipid metabolism^24^. To further investigate the role of PGRN in microglial lipid homeostasis, we generated clonal *Grn* knockout (*Grn*^-/-^) BV2 microglial cell lines using CRISPR/Cas9 (*Grn* KO2 and *Grn* KO8). Depletion of PGRN was confirmed at both protein and mRNA levels by Western blot and quantitative real-time PCR (qRT-PCR), respectively (Fig. 1a-b). As expected, *Grn*^-/-^ BV2 cells showed impaired lysosomal functions, including increased LAMP1 expression and Lysotracker signals, but decreased lysosomal enzyme activities (Fig.S1a-1b), underscoring the critical role of PGRN in lysosomal homeostasis.

**Figure 1.**
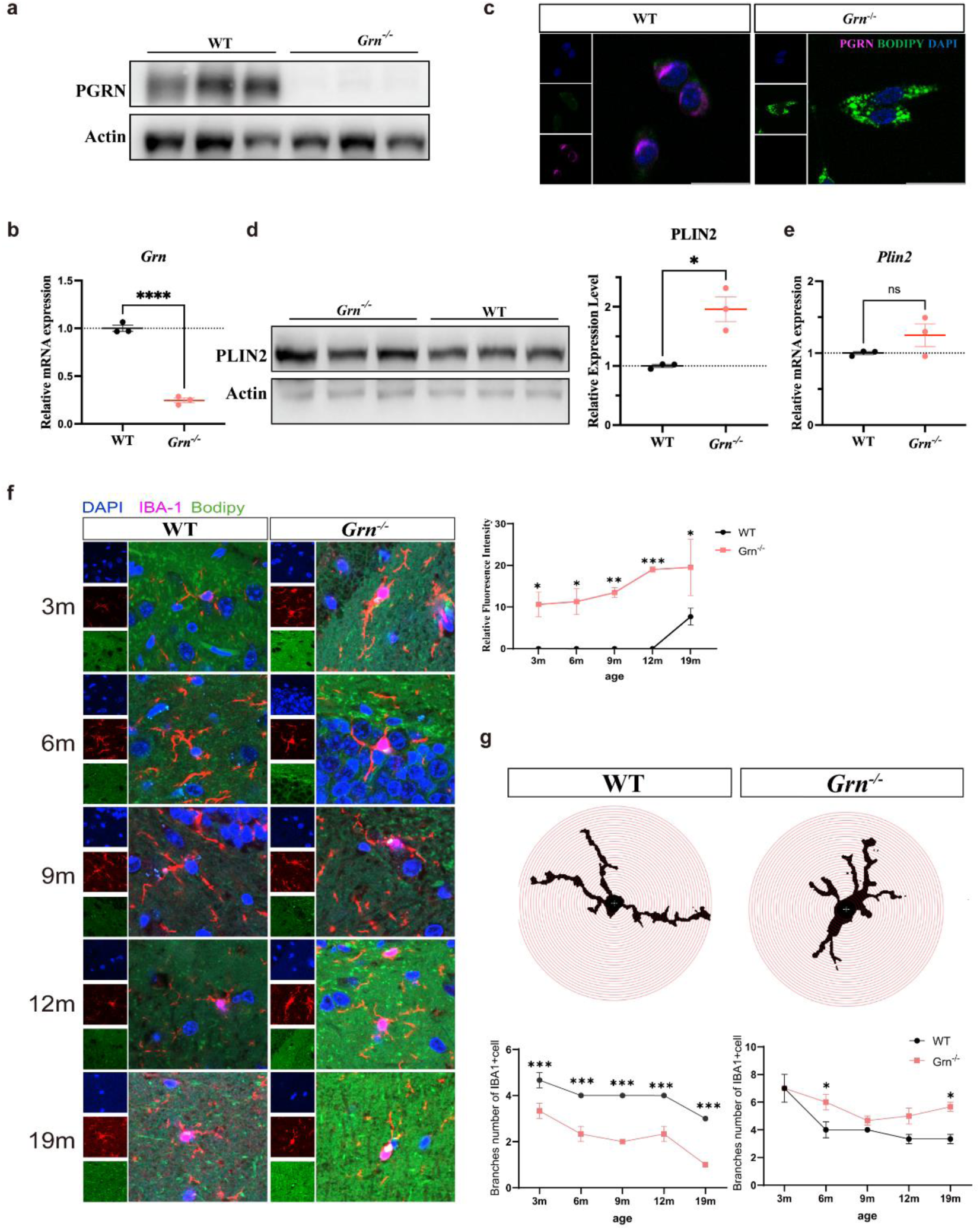
PGRN deficiency drives lipid droplet accumulation in microglia. a. Western blot analysis of PGRN protein in *Grn^-/-^* BV2 microglial cells. b. Analysis of PGRN mRNA levels in *Grn^-/-^* BV2 microglial cells (n=3). c. Representative BODIPY and PGRN fluorescence images in WT and *Grn*^-/-^ BV2 microglial cells. Scale bar, 20 μm. d. Western blot analysis of PLIN2 and statistical results of gray value quantification. In *Grn^-/-^* BV2 microglial cells (n=3). e. Analysis of *PLIN2* mRNA levels in *Grn^-/-^* BV2 microglial cells (n=3). f. Representative BODIPY staining and IBA-1 fluorescence in thalamus of *Grn*^-/-^ mice at different ages and fluorescence quantification of BODIPY (n=3). Scale bar, 20 μm. g. Representative images of Sholl analysis of microglia branches in WT and *Grn*^-/-^ mice. Quantification of the number of proximal branches and distal branches of WT and *Grn*^-/-^ microglia at different ages (n=3).

We next assessed LD accumulation in *Grn*^-/-^ microglia using BODIPY (a specific fluorescent marker for neutral lipids) staining. Immunofluorescence analysis revealed significantly increased formation of spherical LDs within the cytoplasm, accompanied by diminished PGRN expression in *Grn*^-/-^ microglia (Fig. 1c). This finding was independently corroborated by LipidSpot (another neutral lipid-specific dye) staining, which showed intense cytoplasmic red fluorescence in *Grn*^-/-^ microglia (Fig. S2a). Consistent with these observations, protein and mRNA levels of the lipid droplet surface proteins PLIN2 and PLIN3 were markedly elevated in *Grn^-/-^* microglia compared to wild-type (WT) controls (Fig.1d-e, Fig. S2b-2c). These findings collectively suggest that PGRN deficiency induces abnormal LD accumulation in microglia, which is indicative of a LDAM phenotype.

Lipidomic analysis was subsequently conducted to examine the differences of lipid composition between *Grn^-/-^* and WT microglia, focusing on the class I and II lipid metabolites. The analysis further revealed significant alterations in LD composition in *Grn*^-/-^ cells, with a marked increase in glycerolipids (GL) but a decrease in cholesteryl esters (CE) compared to WT microglia (Fig. S2d). These findings suggest that enhanced triglyceride synthesis is the major driver of LD accumulation in *Grn^-/-^* microglia.

To validate LD accumulation caused by PGRN deficiency in vivo, we generated *Grn*^-/-^mice using CRISPR-Cas9 (Fig. S3) and performed immunofluorescence staining on brain sections from mice at different ages. Quantitative analysis of BODIPY staining in Iba1^+^ microglia demonstrated significant LD deposition within thalamic microglia of *Grn*^-/-^ mice (Fig. 1f). Detailed morphological assessment revealed characteristic microglial restructuring: while wild-type microglia maintained small cell bodies with delicate branched processes, *Grn*^-/-^ microglia showed markedly enlarged somata and hyper-ramified processes (Fig.1g). The *Grn*^-/-^ microglia displayed increased proximal and decreased distal branching compared with the wild-type microglia (Fig.1g). These findings collectively indicate that PGRN deficiency induces both microglial activation and pathological LD accumulation in the thalamus in an age-dependent manner.

### Upregulation of GPAT3 in *Grn^-/-^* microglia promotes lipid droplet accumulation

To elucidate the underlying mechanisms of LD accumulation by PGRN deficiency, we performed RNA sequencing (RNA-seq) to analyze transcriptional changes between *Grn*^-/-^ and WT microglial cells. This identified 2,261 differentially expressed genes (DEGs) (p<0.05) in *Grn* KO8, comprising 1,107 upregulated and 1,154 downregulated genes (Fig. 2a, Table S1). RNAseq was also performed in *Grn* KO2 and in a confirmatory assay. Significantly overlapping DEGs were identified across these two clones and the two independent sequencing assays. Concomitantly, *Grn*^-/-^ microglia cell lines exhibited elevated expression of the gene markers related to LD biology^15^ (Fig. S4a-b, Table S1). Notably, Gene Ontology (GO) enrichment analysis of the RNA-seq data revealed that the glycerol-3-phosphate metabolic process was among the most significantly enriched biological processes (BP) (Fig. 2b), suggesting that altered lipid metabolism contribute to the LD accumulation in *Grn*^-/-^ microglia.

**Figure 2.**
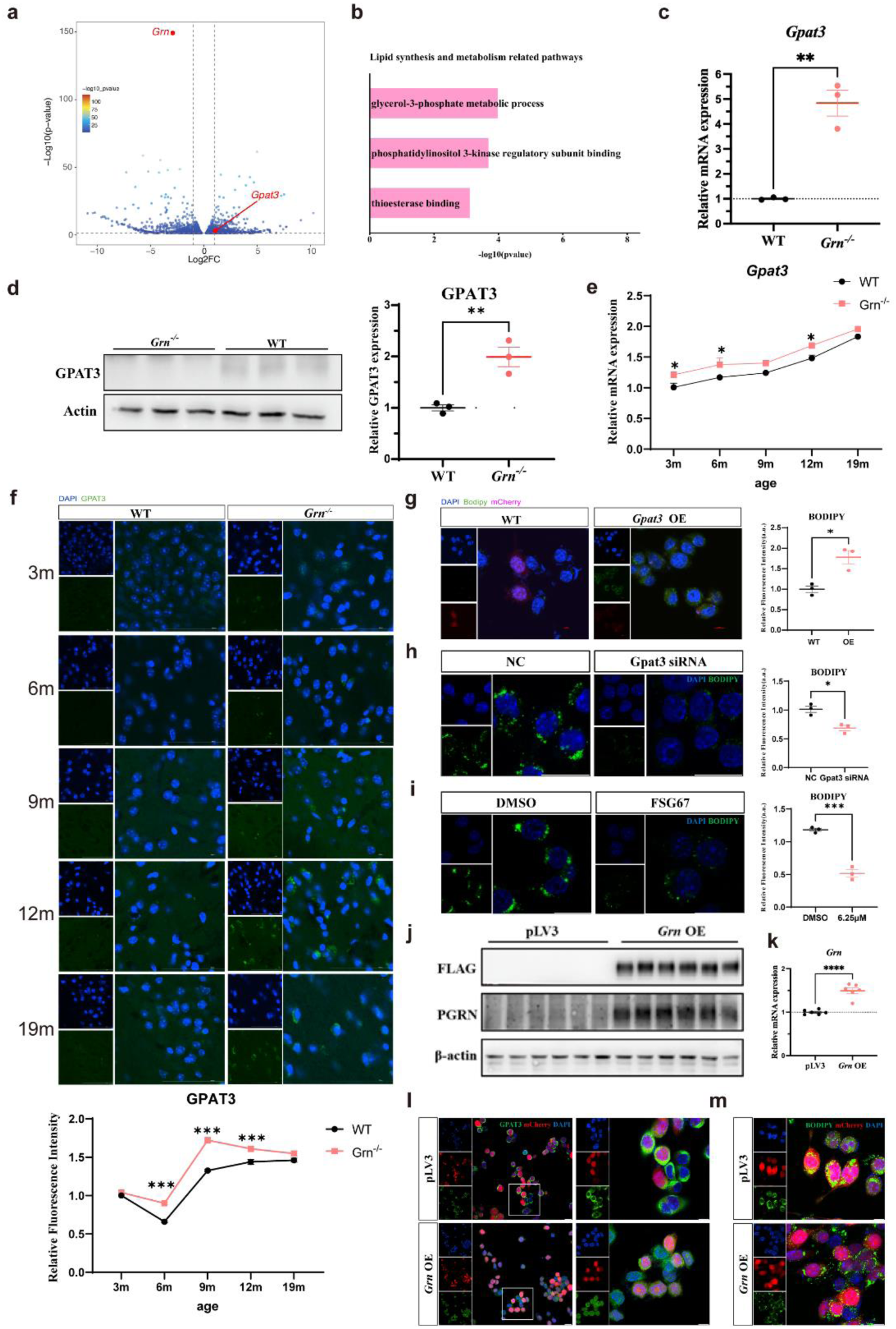
GPAT3 in *Grn^-/-^* microglia promotes lipid droplet accumulation. a. Volcano plot of differentially expressed genes in *Grn^-/-^* BV2 microglial cells. b. GO enrichment analysis of differentially expressed genes (DEGs) from RNA-seq in *Grn*^-/-^ BV2 microglia. c. Analysis of *GPAT3* mRNA levels in *Grn^-/-^* BV2 microglial cells (n=3). d. Western blot analysis of GPAT3 and statistical results of gray value quantification in WT and *Grn^-/-^* BV2 microglial cells (n=3). e. Analysis of *Gpat3* mRNA levels *Grn*^-/-^ mouse brains at different ages (n=3). f. Representative GPAT3 fluorescence in thalamus of *Grn*^-/-^ mice at different ages and fluorescence quantification of GPAT3 (n=3). Scale bar, 20 μm. g. Representative BODIPY staining of GPAT3-overexpressing BV2 microglial cells and fluorescence quantification of BODIPY(n=3). Scale bar, 20 μm. h. Representative BODIPY staining of *Grn*^-/-^ BV2 cells treated with *Gpat3* siRNA or scramble siRNA negative control (NC) and fluorescence quantification of BODIPY signals (n=3). Scale bar, 20 μm. i. Representative BODIPY staining of *Grn*^-/-^ BV2 cells treated with FSG67 or DMSO and fluorescence quantification of BODIPY signals (n=3). Scale bar, 20 μm. j. Western blot analysis of FLAG-tagged PGRN protein in *Grn*^-/-^ BV2 microglial cells after *Grn* restoration. k. Analysis of *Grn* mRNA levels in *Grn*^-/-^ BV2 microglial cells after *Grn* restoration (n=6). l-m. Representative GPAT3 fluorescence and BODIPY staining of *Grn*^-/-^ BV2 microglial cells following PGRN restoration. Scale bar, 20 μm.

Since triglyceride synthesis was predominantly enhanced in *Grn*^-/-^ microglia, we analyzed the expression of GPATs, the critical enzymes for triglyceride biosynthesis, in *Grn*^-/-^ microglia. RNA-seq analysis revealed that GPAT3 but no other GPATs was significantly increased in *Grn^-/-^*microglia compared to WT cells (Fig. 2a, Fig. S4a, Table S1). The upregulation of GPAT3 in *Grn^-/-^* microglia was also confirmed by qRT-PCR and Western blot (Fig. 2c-d). We also analyzed GPAT3 expression in *Grn*^-/-^ mice and found that its expression was upregulated in an age-dependent manner and temporally correlates with the LD formation (Fig. 2e-f). Our results are in line with a previous report showing upregulated expression of GPAT3 in *Grn*^-/-^ microglia^31^. Furthermore, we also observed that microglial GPAT3 expressing increased with aging (Fig. 2e-f), suggesting that GPAT3 may also mediate the LD accumulation upon aging. To confirm a direct link between GPAT3 upregulation and LD formation, we generated GPAT3-overexpressing BV2 microglial cells. BODIPY staining revealed that GPAT3 overexpression was sufficient to increase LD accumulation, recapitulating the observations in *Grn^-/-^* microglia (Fig.2g). Next, to determine whether GPAT3 is functionally required for LD formation in microglia, we silenced *Gpat3* in *Grn*^-/-^ microglia by siRNAs. First, we designed three siRNAs targeting *Gpat3*. After qPCR detection, we found that the knockdown efficiency of siRNA1 was approximately 50%. Therefore, in subsequent experiments, we selected siRNA1 and verified its knockdown efficiency via Western blot (Fig. S5a-b). Knockdown of *Gpat3* could reverse the LD accumulation phenotype (Fig. 2h). Additionally, we pharmacologically inhibited GPATs activity by a small molecule inhibitor FSG67, BODIPY staining revealed that FSG67 treatment led to approximately a 50% reduction in intracellular LDs (Fig. 2i). These results suggest that both siRNA-mediated suppression of GPAT3 expression and pharmacological inhibition of GPAT activity effectively mitigates LD accumulation in *Grn^-/-^* microglia. Therefore, GPAT3 represents a critical regulator of LD formation in microglia.

Next, to determine whether PGRN level regulates GPAT3 expression, we restored PGRN expression (FLAG-tagged) in *Grn^-/-^* cell line (*Grn* OE) using a lentiviral vector co-expressing mCherry. Successful restoration of PGRN expression in *Grn^-/-^* microglia was confirmed by qRT-PCR, Western blot and immunofluorescence staining (Fig.2j-k, Fig. S5c). Importantly, restoring PGRN expression in *Grn*^-/-^ microglia could downregulate GPAT3 level and reduce LD accumulation (Fig.2l-m), as cells positive for mCherry showing significantly decreased GPAT3 signals and reduced BODIPY staining. In addition, qRT-PCR analysis showed that upon PGRN restoration, mRNA levels of the lipid droplet surface protein PLIN2 and PLIN3 decreased (Fig. S5d). Collectively, these results suggest that PGRN deficiency upregulates GPAT3 expression, consequently enhancing triglyceride synthesis and promoting LD formation, whereas PGRN restoration effectively reverses GPAT3 expression and lipid droplet accumulation in *Grn^-/-^* microglial cells.

To determine whether the LD accumulation was due to the impaired lipid degradation. we also analyzed the expression of enzymes for triglyceride degradation. Interestingly, we found that these genes were also upregulated by PGRN depletion (Fig. S5e). Due to the overall increase of triglyceride and LD accumulation, we speculate the upregulation of these metabolic genes was a response rather that a trigger to the LD accumulation.

### PGRN deficiency upregulates GPAT3 via STAT3 pathway

To delineate the mechanisms by which PGRN deficiency upregulates GPAT3, we focused on IL-6/STAT3 pathway, a well-validated, disease-relevant signaling node previously linked to both PGRN deficiency and GPAT3 expression^32,33^. Using complementary in vitro (BV2 microglia) and in vivo (*Grn*^-/-^ mice) models, we demonstrated that PGRN ablation robustly upregulated IL-6 expression (Fig. 3a-c). Concordantly, phosphorylated STAT3 (pSTAT3) - the active, transcriptionally competent form of STAT3 - was significantly elevated in both *Grn*^-/-^ BV2 cells and *Grn*^-/-^ mouse brains (Fig. 3d-e). Critically, PGRN restoration in *Grn*^-/-^ BV2 cells abrogated STAT3 activation (Fig. 3d), confirming PGRN as a direct negative regulator of the IL-6/STAT3 pathway. These results support the hypothesis that PGRN deficiency drives IL-6/STAT3 pathway activation.

**Figure3.**
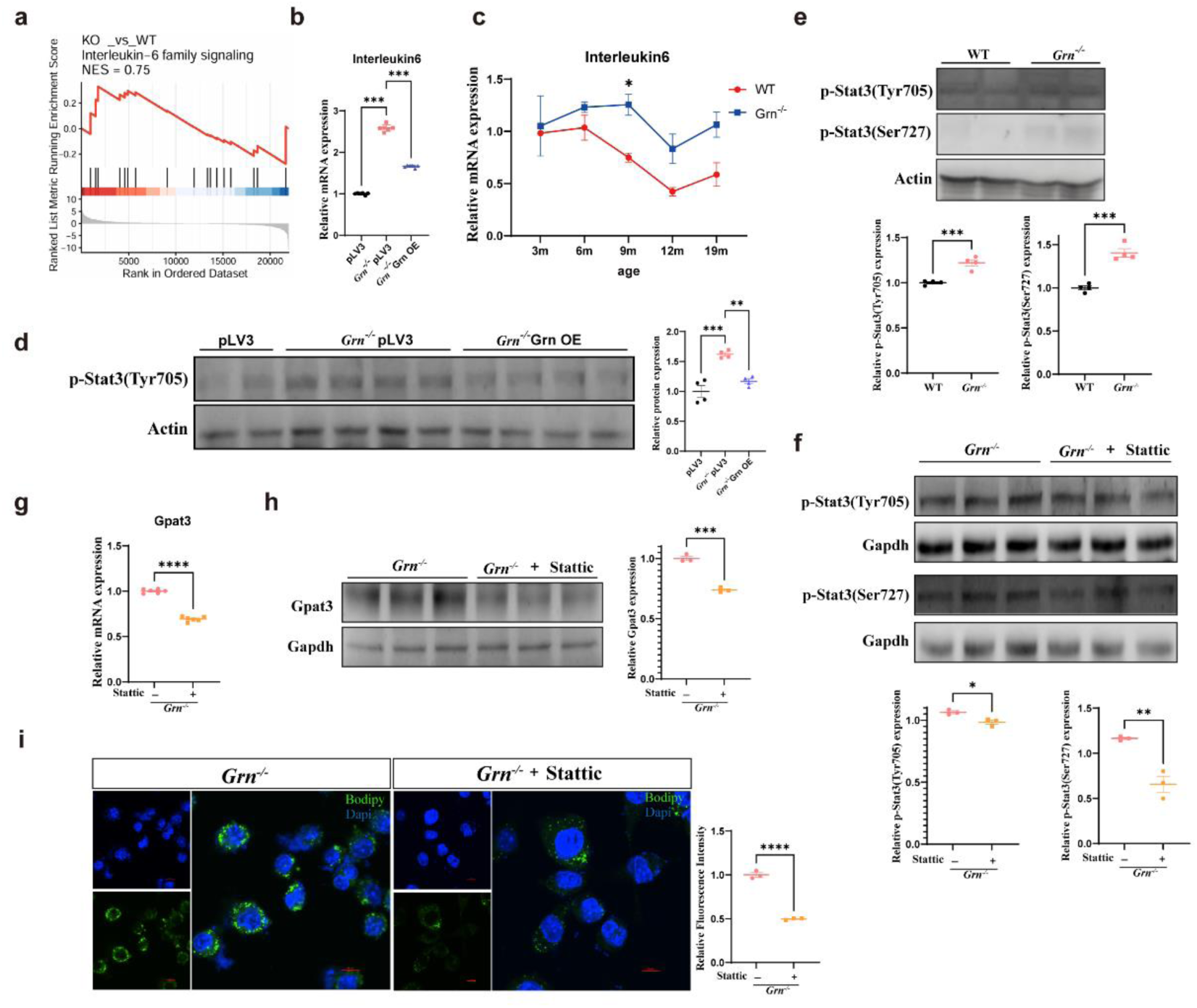
PGRN deficiency upregulates GPAT3 via STAT3 pathway. a. Gene se enrichment analysis (GSEA) of IL-6 signaling in *Grn*^-/-^ BV2 microglial cells. b. Analysis of *IL-6* mRNA levels in *Grn^-/-^* BV2 microglial cells and *Grn*-restored *Grn^-/-^* BV_2_ microglial cells (n=6). c. Analysis of *IL-6* mRNA levels in *Grn^-/-^* mouse brains at different ages (n=3). d. Western blot analysis of p-STAT3(tyr705) and statistical results of gray value quantification. in *Grn^-/-^* BV2 microglial cells and *Grn*-restored *Grn^-/-^* BV2 microglial cells (n=4). e. Western blot analysis of p-STAT3(tyr705) and p-STAT3(ser727) and statistical results of gray value quantification 9-month-old *Grn*^-/-^ mice (n=4). f. Western blot analysis of p-STAT3(tyr705) p-STAT3(ser727) and statistical results of gray value quantification in *Grn^-/-^* BV2 microglial cells and *Grn^-/-^* BV2 microglial cells treated with Stattic (n=3). g. Analysis of *GPAT3* mRNA levels in *Grn^-/-^* BV2 microglial cells and *Grn^-/-^* BV2 microglial cells treated with Stattic (n=6). h. Western blot analysis of GPAT3 and statistical results of gray value quantification in *Grn^-/-^* BV2 microglial cells and *Grn^-/-^* BV2 microglial cells treated with Stattic (n=3). i. Representative BODIPY staining of *Grn^-/-^* BV2 microglial cells and *Grn^-/-^* BV2 microglial cells treated with Stattic and fluorescence quantification of BODIPY (n=3). Scale bar, 20 μm.

To definitively establish STAT3 activation as an upstream driver of GPAT3 upregulation and subsequent LD formation, we employed pharmacological inhibition using Stattic (MCE, HY13818) - a widely used, selective STAT3 inhibitor. Western blot analysis confirmed effective suppression of pSTAT3 levels in Stattic-treated *Grn*^-/-^ BV2 cells (Fig. 3f-g). Functional readouts revealed that STAT3 blockade not only significantly abrogated GPAT3 upregulation induced by PGRN deficiency but also markedly reduced LD accumulation (Fig. 3h-i) - a direct functional consequence of GPAT3 inhibition.

Collectively, these findings mechanistically delineate that PGRN deficiency upregulates GPAT3 and promotes LD formation primarily through the IL-6/STAT3 axis. Beyond advancing basic understanding of PGRN-related neuroinflammation and lipid dysregulation, this work identifies the IL-6/STAT3-GPAT3 cascade as a potential therapeutic target.

### GPAT3-mediated Lipid droplets formation in *Grn*^-/-^ microglia leads to neurotoxicity

To dissect the neurotoxic potential of LDAM, we collected microglial conditioned media (MCM) from WT and *Grn^-/-^* microglia and treated N2a neuronal cells with these MCM for 48 hours to assess cytotoxicity. CCK-8 assays revealed that *Grn^-/-^* MCM reduced N2a cell viability to approximately 80% of WT MCM controls (Fig. 4a), a statistically significant reduction that mirrors the microglial dysregulation observed in PGRN-deficient neurodegenerative diseases such as FTD. Concordantly, lactate dehydrogenase (LDH) release -an established maker of neuronal membrane damage - was significantly elevated by 1.4-fold in the *Grn^-/-^* MCM-treated cells (Fig. 4b), confirming that conditioned media from PGRN-deficient microglia induced considerable neuronal damage. Notably, DCFH-DA assay demonstrated markedly higher reactive oxygen species (ROS) levels in the *Grn^-/-^* MCM-treated cells compared to WT MCM controls (Fig. 4c), suggesting a critical mechanistic link between microglial PGRN deficiency, oxidative stress, and neuronal damage and filling a key gap in understanding LDAM-mediated neurotoxicity.

**Figure 4.**
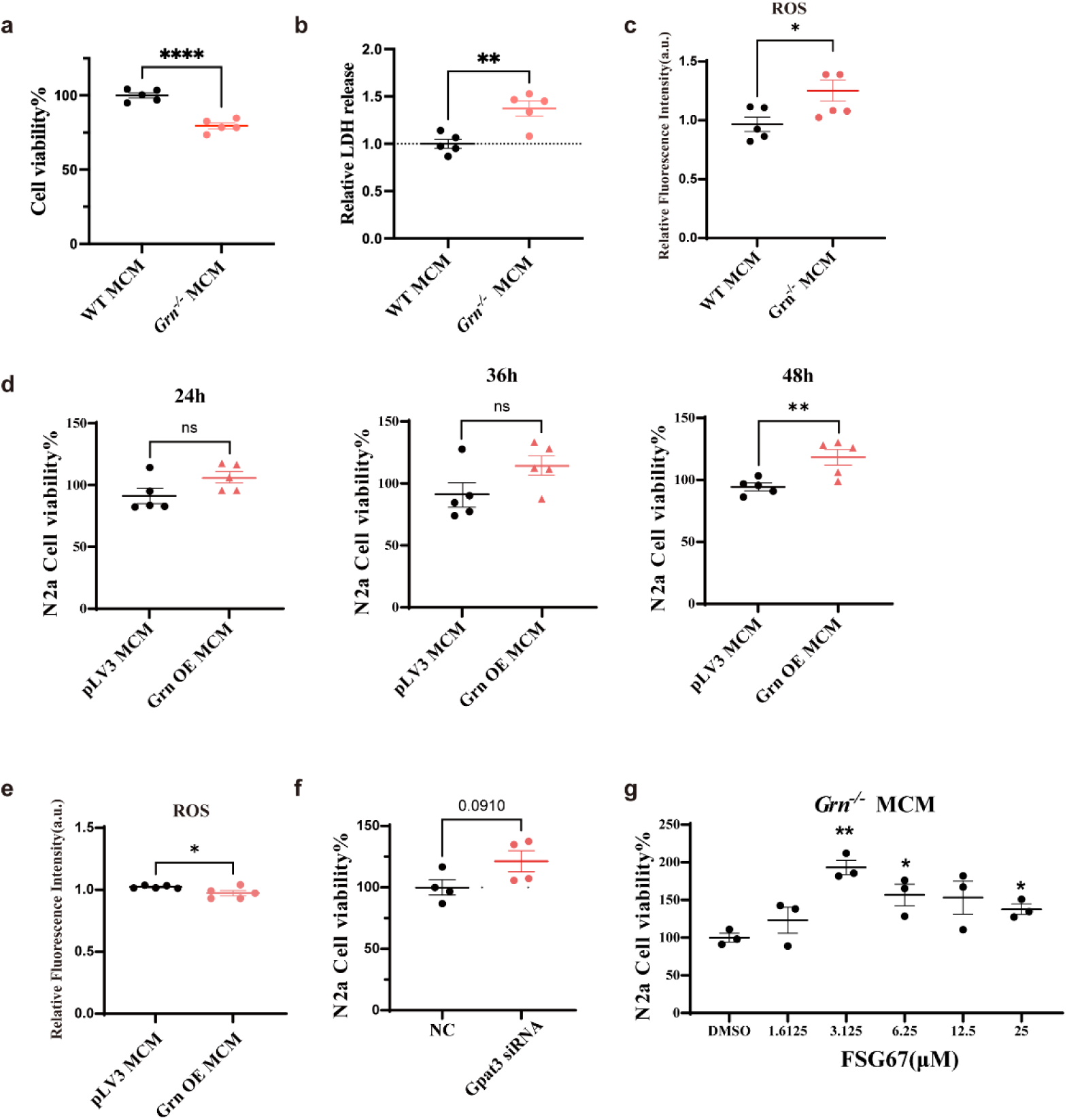
GPAT3-mediated Lipid droplets formation in *Grn*^-/-^ microglia leads to neurotoxicity. a. Cell viability assay of mouse neuronal N2a cells treated with conditioned media from either BV_2_ microglia cells or *Grn*^-/-^ BV2 microglial cells (n = 5). b. Quantitative analysis of intracellular lactate dehydrogenase (LDH) levels in N2a cells treated as in (A) (n = 5). c. Reactive oxygen species (ROS) levels in N2a cells treated as in (A) (n = 5). d. Cell viability assay of mouse neuronal N2a cells treated with conditioned media from either *Grn*^-/-^ microglial cells or *Grn*-rescued *Grn*^-/-^ microglia (n = 5). e. Intracellular reactive oxygen species (ROS) levels in mouse neurons treated as in (D) (n = 5). f. N2a cell viability after treatment with conditioned media from either *Grn*^-/-^ microglial cells or siRNA-*Gpat3* treated *Grn*^-/-^ microglial cells (n = 4). g. N2a cell viability after treatment with conditioned media from either *Grn*^-/-^ microglial cells or FSG67-treated *Grn*^-/-^ microglial cells (n = 3).

To validate PGRN as a therapeutic modulator, we next treated N2a cells with MCM from *Grn^-/-^* microglial cells transfected with either empty pLV3 vector (control) or pLV3-*Grn* (*Grn* OE) for 24, 36, and 48 hours. CCK-8 assay revealed a time-dependent rescue effect. N2a cells treated with *Grn* OE MCM after 48 hours exhibited an increase in viability compared to those treated with vector control MCM (Fig. 4d). ROS measurement by DCFH-DA assay further confirmed that *Grn* OE MCM significantly reduced ROS levels in N2a cells compared to vector control MCM (Fig. 4e). These results suggest that restoration of PGRN expression in *Grn^-/-^*microglia could mitigate their neurotoxicity to N2a cells. These findings not only confirm that PGRN deficiency is causative of LDAM neurotoxicity but also provide proof-of-concept for PGRN-based therapeutic strategies.

We finally interrogated whether the GPAT3-mediated LD accumulation directly drives neurotoxicity. N2a cells were treated with MCM from *Grn^-/-^* BV2 cells subjected to *Gpat3* siRNA or scramble siRNA (negative control). Four independent CCK-8 assays consistently showed a trend toward increased N2a viability in the *Gpat3* siRNA-conditioned medium (siMCM) compared to the negative control (NC MCM) (p = 0.0910, Fig. 4f). Additionally, pharmacological inhibition of GPAT3 with FSG67 dose-dependently enhanced N2a cell viability in response to *Grn^-/-^* MCM (Fig. 4g). These results indicate that both genetic and pharmacological targeting of GPAT3 effectively reverses LD accumulation and significantly attenuates LDAM-mediated neurotoxicity, highlighting GPAT3 as a tractable therapeutic target for LDAM-associated neurodegeneration.

### Restoration of PGRN in microglia or inhibition of GPAT3 activity rescues neuropathology and behavioral phenotypes of *Grn*^-/-^ mice

To translate our in vitro mechanistic insights into a clinically relevant preclinical model, we next interrogated whether targeting PGRN-GPAT3 axis ameliorates the neuropathological and behavioral deficits in *Grn*^-/-^ mice - an established model of PGRN deficiency-related neurodegeneration such as FTD. Leveraging stereotaxic delivery of adeno-associated virus (AAV), we selectively expressed PGRN in thalamic microglia of 8.5-month-old *Grn*^-/-^ mice via bilaterally stereotaxic injection of AAV-MG1.2-mCx3cr1-mGrn-eGFP (MG1.2 serotype and Cx3cr1 promoter ensures microglia-specific expression, AAV-*Grn*; Control vector AAV-MG1.2-mCx3cr1-eGFP: AAV-CTRL). Brain tissues were harvested 4 weeks after injection. Confocal immunofluorescent imaging revealed that microglial PGRN restoration significantly reduced GPAT3 expression; concurrent BODIPY staining confirmed a marked attenuation of LD accumulation in *Grn*^-/-^ microglia (Fig. 5a-b). Critically, immunostaining for IBA1 (a microglial activation marker) and GFAP (an astrocytes reaction marker) revealed specific GFP signals in microglia but not in astrocytes, thereby confirming the cell-type specificity of the AAV-mediated expression (Fig. 5c-d). Furthermore, these stainings also demonstrated a robust attenuation of neuroinflammation following PGRN restoration (Fig. 5c-d). These findings validate the PGRN-GPAT3-LD axis as a conserved pathogenic driver in vivo and highlight AAV-mediated microglia-specific gene therapy as a precision translational strategy.

**Figure 5.**
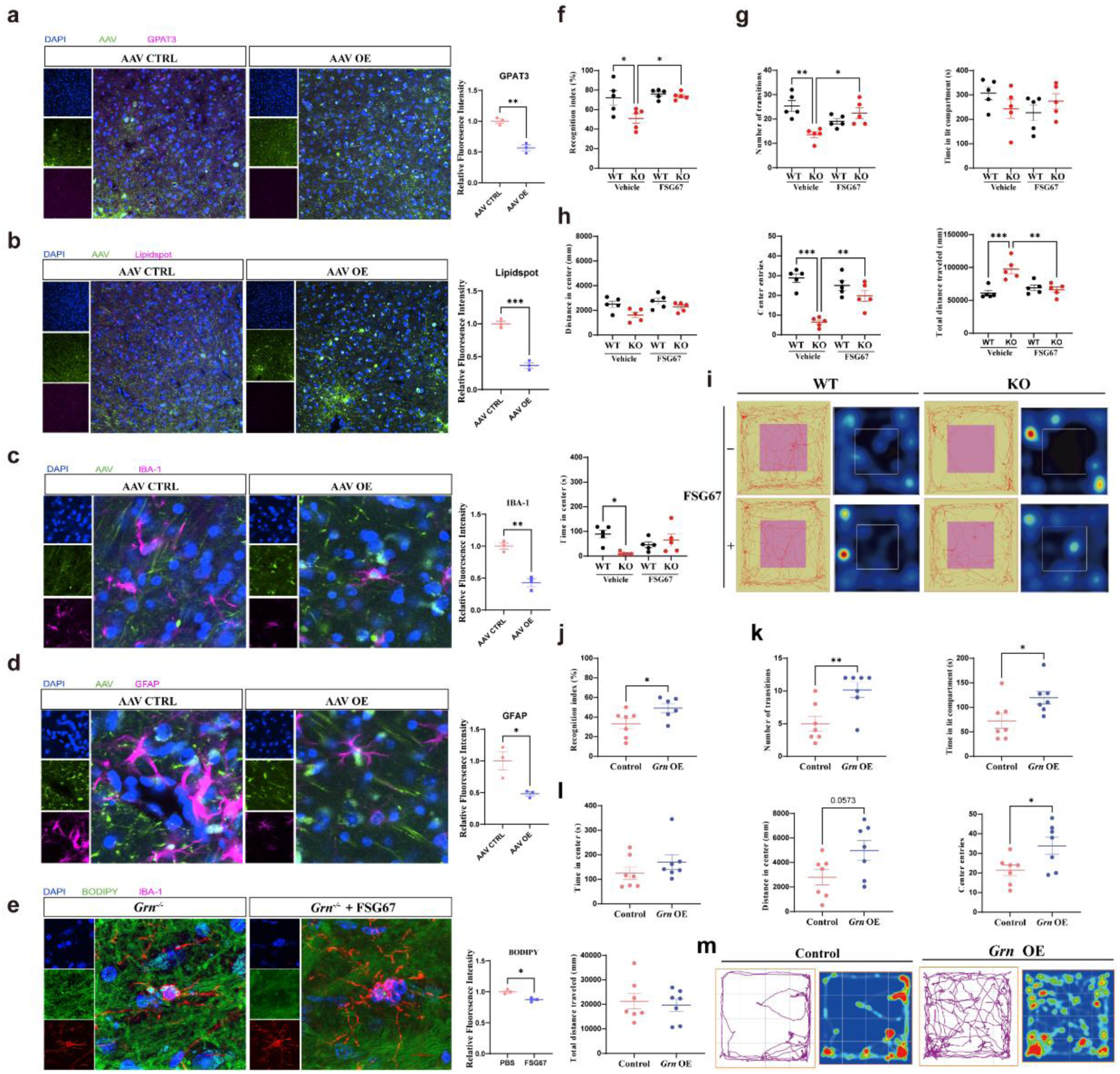
Restoration of PGRN in microglia or inhibition of GPAT3 activity ameliorates neuropathology and behavioral phenotypes of *Grn*^-/-^ mice. a. Representative GPAT3 fluorescence in thalamus of *Grn*^-/-^ mice injected with AAV-CTRL or AAV-*Grn* and fluorescence quantification of GPAT3 signals (n = 3). Scale bar, 20 μm. b. Representative LipidSpot staining in thalamus of *Grn*^-/-^ mice injected with AAV-CTRL or AAV-*Grn* and fluorescence quantification of LipidSpot (n = 3). Scale bar, 20 μm. c. Representative IBA1 fluorescence in thalamus of *Grn*^-/-^ mice injected with AAV-CTRL or AAV-*Grn* and fluorescence quantification of IBA1 signals (n = 3). Scale bar, 20 μm. d. Representative GFAP fluorescence in thalamus of *Grn*^-/-^ mice injected with AAV-CTRL or AAV-*Grn* and fluorescence quantification of GFAP signals (n = 3). Scale bar, 20 μm. e. Representative BODIPY and IBA1 fluorescence staining in thalamus of *Grn*^-/-^ mice treated with FSG67 or vehicle and fluorescence quantification of BODIPY in IBA^+^ cells (n = 3). Scale bar, 20 μm. f. Recognition working memory assessed by the novel object recognition test in WT and *Grn*^-/-^ mice treated with FSG67 or vehicle (n = 5 for each group). g. Anxiety behavior assessed by the light-dark transition test in WT and *Grn*^-/-^ mice treated with FSG67 or vehicle (n = 5 for each group). h. Anxiety behavior assessed by the open field test in WT and *Grn*^-/-^ mice treated with FSG67 or vehicle (n = 5 for each group). i. Representative open-field trajectory heatmaps for the groups in Fig. 5h (SuperMaze software). j. Recognition working memory assessed by the novel object recognition test in *Grn*^-/-^ mice injected with AAV-CTRL or AAV-*Grn* (n = 7 for each group). k. Anxiety behavior assessed by the light-dark transition test in *Grn*^-/-^ mice injected with AAV-CTRL or AAV-*Grn* (n = 7 for each group). l. Anxiety behavior assessed by the open field test in *Grn*^-/-^ mice injected with AAV-CTRL or AAV-*Grn* (n = 7 for each group). m. Representative open-field trajectory heatmaps for the groups in Fig. 5l (AnyMaze software).

To further advance translational potential, we evaluated the therapeutic efficacy of FSG67 via intraperitoneal injection in 8.5-month-old *Grn*^-/-^ mice. Indeed, 4 weeks after injection, FSG67 administration significantly suppressed GPAT3 enzymatic activity, as evidenced by a profound reduction in LD accumulation in *Grn*^-/-^ mice (Fig. 5e). More importantly, behavioral assessments including novel object recognition test, light-dark transition test and open field test revealed that FSG67-mediated GPAT3 inhibition significantly improved the behavioral deficits in *Grn*^-/-^ mice compared to vehicle controls (Fig. 5f-i). Parallel behavioral tests confirmed that microglial PGRN restoration also rescued the behavioral phenotype of *Grn*^-/-^ mice (Fig. 5j-m), corroborating that targeting the PGRN-GPAT3 axis - via either gene replacement or small-molecule inhibition - ameliorates disease-relevant deficits.

Collectively, these in vivo findings establish the PGRN-GPAT3 axis as a tractable, translationally viable therapeutic target for PGRN deficiency-related neuropathology and neurodegeneration. By bridging in vitro mechanistic discoveries to preclinical efficacy in a disease-relevant mouse model, our work provides a critical “bench-to-bedside” proof-of-concept: (1) AAV-mediated microglia-specific PGRN restoration represents a precision gene therapy strategy; (2) FSG67-based GPAT3 inhibition offers a readily translatable small-molecule approach. This dual-pronged therapeutic paradigm addresses an unmet clinical need for FTD and other PGRN-associated neurodegenerative disorders, underscoring the high impact of our findings for both basic neuroscience and clinical translation.

## Discussion

PGRN deficiency is strongly associated with various neurodegenerative disorders, most notably frontotemporal dementia (FTD-*GRN*), yet the molecular cascades linking PGRN loss to neuronal dysfunction and death remain incompletely resolved^34^. In this study, we identify a novel pathogenic pathway wherein PGRN deficiency drives LD accumulation in microglia through the STAT3-GPAT3 axis, ultimately eliciting neurotoxicity and behavioral impairments. These findings unravel a critical role for microglial metabolic reprogramming in PGRN-related neuropathology and highlight actionable therapeutic targets for FTD-*GRN* and potentially other age-related neurodegenerative conditions.

Our work first confirms that PGRN deficiency disrupts microglial lipid homeostasis, leading to pathological LD accumulation. Consistent with prior evidence implicating PGRN in lipid metabolism^24^, we demonstrate robust LD formation in *Grn*^-/-^ microglia-both in vitro (BV2 cells) and in vivo (mouse brain) - as validated by BODIPY/LipidSpot fluorescent staining and upregulated expression of LD surface markers PLIN2 and PLIN3 at both mRNA and protein levels. Lipidomic profiling further delineates that this accumulation is predominantly driven by enhanced glycerolipid (GL) biosynthesis, particularly triglyceride synthesis, rather than impaired lipid degradation. This conclusion is supported by two key observations: (1) selective upregulation of triglyceride biosynthetic enzymes (most notably GPAT3) in *Grn*^-/-^microglia; and (2) failure of upregulated lipid degradation-related genes to rescue LD accumulation. Notably, the age-dependent progression of LD deposition in the thalamus of *Grn*^-/-^ mice mirrors the progressive neurodegenerative phenotype of FTD-*GRN*, suggesting that microglial LD accumulation may serve as a temporal driver of disease progression.

A central novel finding of this study is the identification of GPAT3 as a critical mediator of LD accumulation in PGRN-deficient microglia. RNA-sequencing and functional validation experiments collectively demonstrate that GPAT3 upregulation is both necessary and sufficient for LD formation: (1) GPAT3 overexpression in wild-type (WT) BV2 cells recapitulates the *Grn*^-/-^ LD phenotype; (2) siRNA-mediated GPAT3 knockdown reverses LD accumulation in *Grn*^-/-^ microglia; and (3) pharmacological inhibition of GPAT activity (via FSG67) reduces LD formation by ∼50% in *Grn*^-/-^ cells. These results extend a prior report of GPAT3 upregulation in *Grn*^-/-^ microglia by establishing GPAT3 as a functional mediator of lipid dyshomeostasis. Furthermore, the age-dependent upregulation of GPAT3 in *Grn*^-/-^ mice and aged WT mice suggests that GPAT3 may act as a common mediator of LD accumulation in both genetic (PGRN deficiency) and age-related microglial dysfunction - an important implication given the heightened risk of neurodegeneration with aging. Future studies should prioritize validating GPAT3 upregulation in microglia isolated from FTD-*GRN* patient brain tissues, which would directly strengthen its clinical relevance.

We further dissect the upstream signaling mechanism by which PGRN deficiency regulates GPAT3 expression: PGRN ablation induces IL-6 production in microglia, leading to phosphorylation and activation of STAT3 (pSTAT3), which subsequently transactivates GPAT3. This is in line with a previous observation on hepatocellular carcinoma cells^33^. Functional validation confirms that blocking pSTAT3 significantly attenuates both GPAT3 upregulation and LD formation in *Grn*^-/-^ microglia, establishing the STAT3 signaling as a key regulatory pathway. This finding integrates two well-characterized features of PGRN deficiency - neuroinflammation and metabolic dysregulation - by demonstrating crosstalk between inflammatory signaling and lipid metabolism in microglia. Importantly, this crosstalk suggests that anti-inflammatory strategies targeting the IL-6/STAT3 axis may simultaneously normalize lipid homeostasis, offering a dual-benefit therapeutic approach. Interestingly, the impact of PGRN on STAT3 signaling in microglia differs from what has been reported in adipocytes^32^, implying a cell type-specific regulatory mechanism.

Critically, we demonstrate that GPAT3-mediated LD accumulation endows *Grn*^-/-^microglia with neurotoxic properties. Conditioned media (MCM) from *Grn*^-/-^ microglia reduces N2a neuronal viability, increases lactate dehydrogenase (LDH) release, and elevates reactive oxygen species (ROS) production in neurons - all hallmarks of neuronal damage. These neurotoxic effects are directly linked to GPAT3-mediated LD accumulation, as evidenced by their reversal following: (1) GPAT3 knockdown; (2) GPAT pharmacological inhibition; or (3) PGRN restoration in *Grn*^-/-^ microglia. These data support the concept that LDAM represent a distinct neurotoxic phenotype, distinguishable from both homeostatic and classical pro-inflammatory microglia^15^. The increased ROS production in neurons treated with *Grn*^-/-^ MCM suggests oxidative stress as a key mediator of neurotoxicity, which may arise from either impaired LD metabolism in microglia or altered secretion of cytotoxic factors. Importantly, in vivo validation confirms that targeting the PGRN-GPAT3 axis - either via microglia-specific PGRN restoration (AAV-mediated) or systemic GPAT inhibition (FSG67) - not only reverses LD accumulation and neuroinflammation but also ameliorates behavioral deficits in *Grn*^-/-^ mice. These preclinical findings strongly validate the PGRN-GPAT3 axis as a therapeutic target, highlighting the potential of GPAT3 inhibitors or PGRN-based replacement therapies for FTD-GRN.

Several limitations of this study should be acknowledged to guide future research. First, while the IL-6/STAT3 pathway is identified as a key regulator of GPAT3, other signaling cascades may contribute to LD accumulation in PGRN-deficient microglia, warranting comprehensive investigation. Second, although our PGRN restoration experiments utilized microglia-specific AAV vectors and stereotaxic injection to achieve targeted delivery, the systemic administration of FSG67 may still lead to off-target effects in peripheral tissues. Future studies could explore brain-penetrant, microglia-selective GPAT3 inhibitors to enhance therapeutic specificity and minimize systemic side effects. Third, the reliance on BV2 microglial cells and mouse models necessitates validation in primary human microglia and FTD-*GRN* patient brain tissues to confirm clinical applicability. Finally, the precise molecular mechanisms by which LD accumulation induces neurotoxicity (e.g., secretion of specific cytokines/chemokines, lipid mediators, or direct microglia-neuron synaptic interactions) remain to be fully elucidated.

In conclusion, our study delineates a novel PGRN-STAT3-GPAT3-LD pathway that contributes to neurotoxicity and neurodegeneration in PGRN deficiency. These findings establish GPAT3 as a critical mediator of microglial lipid metabolic dysregulation and highlight the therapeutic potential of targeting the PGRN-GPAT3 axis for FTD-*GRN*. By linking PGRN loss to the convergence of microglial metabolic reprogramming and neuroinflammation, our work provides a new framework for understanding PGRN-related neurodegeneration and offers actionable targets for preclinical drug development.

## Materials and Methods

### Ethical statement

All mice were housed in the specific-pathogen-free animal facility of Fudan University with ad libitum access to food and water. Housing conditions were maintained at 22 °C, 45–65% relative humidity, and a 12 h-12 h light–dark cycle. All of the animal experiment procedures were approved by ethics committee of Department of Laboratory Animals, Fudan University.

### Generation and genotyping of *Grn* knockout mice

To generate *Grn* knockout mice, exons 2-13 of the *Grn* coding region were deleted using CRISPR/Cas9 gene editing. The targeting construct was microinjected into pronuclei of fertilized C57BL/6J oocytes to produce F0 founder mice. Germline-transmitted F0 mice were identified and outcrossed with wild-type C57BL/6J mice to obtain F1 heterozygotes. F1 heterozygotes were then intercrossed to generate F2 progeny, including *Grn^-/-^* homozygous mutants and wild-type littermate controls. For genotyping, genomic DNA was isolated from tail biopsies and analyzed by PCR using two primer sets: Forward primers 1/Reverse primers 1 (detecting the knockout allele) and Forward primers 2/ Reverse primers 2 (detecting the wild-type allele).

For genotype determination: 1) Amplification with Forward primer 1/ Reverse primer 1 produced a 474-bp band if the target gene was successfully deleted, 2) Amplification with Forward primer 2/ Reverse primer 2 produced a 732-bp band if the wild-type allele remained intact. Forward primer 1: 5’-AAG GGC TCA CCA TGA AGT TAG C-3’; Reverse primer 1: 5’-AAC CGG AAG AAA TGG CAG TTT G-3’; Forward primer 2: 5’-AGG ATG TCG ATT TTA TCC AGC CTC-3’; Reverse primer 2: 5’-TGG CGC AAC AAA TCA GAT ACT ATT C-3’.

### Construction of *Grn* knockout cell lines

Mouse BV2 microglial cells were cultured at 37°C in a humidified atmosphere with 5% CO₂ in DMEM supplemented with 10% (v/v) Fetal bovine serum (FBS) and 1% (v/v) penicillin–streptomycin–Glutamax (MeilunBio). Human embryonic kidney 293T cells were maintained at 37°C with 5% CO₂ in DMEM supplemented with 10% (v/v) FCS and 1% (v/v) penicillin–streptomycin. All cell lines used in this study were purchased from the American Type Culture Collection (ATCC).

The sgRNAs targeting the mouse *Grn* gene were designed using the CRISPR design tool CRISPOR (http://crispor.tefor.net/). After synthesis, the sgRNA oligonucleotides (Forward primer: 5’-CAC CGC TGG CTG GCC TTC GCG GCA GT-3’, Reverse primer: 5’-TAA AAC TGC CGC GAA GGC CAG CCA GC-3’) annealed and cloned into the LentiV2 CRISPR/Cas9 vector. Successful plasmid construction was confirmed by Sanger sequencing (conducted by Sangon Biotech, Shanghai). The sequencing results were aligned with the reference plasmid sequence using SnapGene software for verification.

Twenty-four hours before transfection, 293T cells were seeded in 10-cm culture dishes to reach 70-90% confluency at the time of transfection. Upon cell attachment, a transfection mixture was prepared by mixing 4 μg of PMD2.G, 3 μg of pSPAX2, and 6 μg of the recombinant plasmid in 1 mL of DMEM medium. After incubating the mixture at room temperature for 5 minutes, 26 μL of HighGene transfection reagent (Abcloanl, RM09014P) was added and mixed thoroughly. The resulting plasmid-HighGene complex was then evenly distributed to the 293T cells. Following 4-6 hours of transfection, half of the culture medium was replaced with fresh complete medium.

The viral supernatant was harvested 48 hours post-transfection and filtered through a 0.45 μm sterile filter (Millipore, HAWP04700). The clarified viral suspension was then applied to BV_2_ microglial cells that had been pre-seeded in 6-well plates. Following 48-72 hours of viral transduction, the culture medium was replaced with complete medium containing puromycin (1-2 μg/mL, concentration to be optimized) for selection over a 2-day period. The resulting stable *Grn*-knockout BV_2_ cell lines were subsequently validated by both PCR genotyping and Western blot analysis.

### Generation of *Grn* or *Gpat3* overexpression cells lines

The *Grn* overexpression vector was constructed using a double restriction enzyme digestion method, with NheI and BamHI (NEB, R3131S and R3136S) selected as the upstream and downstream restriction sites, respectively. *Grn*-specific primers (Forward primer: 5’-CTA GCT AGC ATG TGG GTC CTG ATG AG-3’, Reverse primer: 5’-CCG GAA TTC TTA CAG TAG CGG TCT TGG-3’) were designed to amplify the mouse *Grn* coding sequences using cDNA templates derived from mouse microglial cells. The PCR-amplified *Grn* fragment was purified by gel extraction and subsequently ligated into the linearized overexpression vector. Finally, the ligation product was submitted to Sangon Biotech (Shanghai) for sequence verification.

To establish *Grn* overexpressing cells, the *Grn* overexpression plasmid was packaged into lentiviral particles and transduced into *Grn*^-/-^ microglial cells. Viral transduction was performed following the protocol detailed in the “Construction of *Grn* knockout plasmid and generation of *Grn*^-/-^ microglial cells” section. Stable transformants were selected using G418 (Geneticin; concentration range 200-1000 μg/mL, optimized based on cell line sensitivity) for approximately 10 days. The resulting *Grn* overexpressing stable cell line was validated by both PCR amplification and Western blot analysis. Similar strategy was used to establish *Gpat3*-overexpressing cells. *Gpat3*-specific primers (Forward primer: 5’-CTA GCT AGC GCC ACC ATG GAG GGC GCA GAC CTG G-3’, Reverse primer: 5’-CGC GGA TCC GTC CCT CGC CAG GTT GGG AGA T-3’) were used to amplify its coding sequences.

### Protocol for simultaneous RNA and protein extraction from mouse brain tissue

Mice were anesthetized via intraperitoneal injection of sodium pentobarbital (50 mg/kg body weight) and underwent transcardial perfusion initiated through the left ventricular apex with simultaneous right atrial incision. Vascular flushing was performed with 20–30 mL sterile physiological saline (5–10 mL/min) until hepatic effluent cleared, followed by fixation with 100–200 mL of ice-cold 4% PFA at the same flow rate. After perfusion, brains were rapidly extracted and sagittally sectioned along the midline using a stainless-steel brain matrix, with hemispheres allocated for parallel biochemical and histochemical studies.

For biochemical analyses, tissue hemispheres were weighed (wet weight recorded to ±0.1 mg precision) and homogenized in 1.5 mL of hypotonic 0.25 M sucrose solution containing 1× protease/phosphatase inhibitors, using a 3 mm diameter stainless steel bead in a pre-chilled tissue homogenizer (2 × 60 sec pulses at 6 m/sec, with 30 sec ice intervals). The homogenate was centrifuged at 3,000 ×g (4°C, 3 min) to generate a post-nuclear supernatant (PNS) fraction.

Protein extracts were prepared by supplementing 500 μL PNS with Complete Mini EDTA-free protease inhibitor cocktail (Roche, 1 tablet/10 mL), aliquoted, and flash-frozen in liquid nitrogen before storage at −80°C. For concurrent RNA isolation, 250 μL PNS was vortex-mixed with an equal volume of QIAzol lysis reagent (Qiagen), incubated at room temperature (5 min), and centrifuged (15,000 × g, 5 min, 4°C). The aqueous phase was combined with molecular-grade ethanol, incubated (−20°C, 30 min), and processed through RNeasy Mini spin columns (Qiagen) per manufacturer’s instructions. RNA integrity was verified by NanoDrop 3000 spectrophotometry (Thermo Fisher) with acceptance thresholds of A260/A280 ≥1.8 and A260/A230 ≥2.0, with all procedures conducted in an RNase-free workstation using DEPC-treated pipette tips and microtubes.

### Western blot analysis

Cellular or tissue protein samples were lysed in RIPA buffer supplemented with protease inhibitor cocktail (MeilunBio, MB12707-1) on ice for 15 minutes. Following thorough vortex mixing, the lysates were centrifuged at 12,000 × g (4°C) for 15 minutes to obtain clarified supernatants containing total protein extracts. Protein concentrations were quantified using the BCA Protein Assay Kit (MeilunBio, MA0082-1), after which samples were prepared in 4× Laemmli buffer at a final concentration of 20 μg protein per 15 μL and heat-denatured at 95°C for 5 minutes. The denatured samples were then loaded onto SDS-polyacrylamide gels, and electrophoresis was performed initially at 80 V until dye front migration, then increased to 120 V. For protein transfer, resolved proteins were electrophoretically blotted onto nitrocellulose membranes at 300 mA constant current for 1 hour at room temperature. Membranes were subsequently blocked with 5% non-fat milk in TBST (10 mM Tris-HCl, 150 mM NaCl, 0.1% Tween-20, pH 7.6) for 1 hour at room temperature with constant agitation. Primary antibody incubation was carried out overnight at 4°C with gentle shaking using antibodies diluted in blocking buffer, followed by three 5-minute TBST washes under agitation. Membranes were then incubated with species-matched HRP-conjugated secondary antibodies (diluted 1:5,000-1:10,000 in blocking buffer) for 1 hour at room temperature with shaking, followed by three additional 5-minute TBST washes. For signal detection, Tanon™ Femto-sig ECL (Tanon, 180-506) was prepared by 1:1 mixing of oxidation and luminol solutions and applied uniformly to the membrane surface. Protein bands were visualized and quantified using the TIANGEN Imaging System. Antibodies used in this study are: PGRN (R&D, AF2557, 1:1,000), LAMP1 (Abcam, ab208943, 1:1,000), GPAT3 (Proteintech, 20603-1-AP, 1:2,000), PLIN2 (Abclonal, A24464, 1:1,000), PLIN3 (Abclonal, A1050, 1:1,000), pSTAT3(Ser727) (Proteintech, 60479-1-Ig, 1:1,000), pSTAT3(Tyr705) (Beyotime, AF5941, 1:1,000), FLAG tag (Abclonal, AE005, 1:1,000), β-tubulin (Abmart, M20005), β-actin (Proteintech, 66009-1-Ig), GAPDH (Millipore, MAB374), HRP-conjugated Goat Anti-Rabbit IgG (H+L) (Proteintech, SA00001-2, 1:5,000), HRP-conjugated Goat Anti-Mouse IgG (H+L) (Proteintech, SA00001-1, 1:50,000).

### Quantitative Real-Time Polymerase Chain Reaction (qRT-PCR)

Total RNA was isolated from cellular or tissue samples using the EZ-press RNA Purification Kit (EZB, B0004D), followed by first-strand cDNA synthesis performed with the 4× EZscript Reverse Transcription Mix II (EZB-RT2) according to the manufacturer’s protocols. The synthesized cDNA products were diluted 1:4 with nuclease-free water prior to PCR amplification. Quantitative real-time PCR (qRT-PCR) reaction mixtures were prepared in triplicate for each biological sample and target gene combination, then dispensed into a 384-well optical reaction plate. Following sample loading, the plate was sealed with optically clear adhesive film, briefly centrifuged at 1000 × g for 1 minute to ensure proper mixing and reaction settlement, and subsequently analyzed using a Roche LC480 Real-Time PCR System under standard cycling conditions. Primer sequences: *Grn*, Forward primer: 5’-GTG TTG TGA GGA TCA CAT TC-3’, Reverse primer: 5’-CTA TGA CCT TCT TCA TCC AG-3’; *Lamp1*, Forward primer: 5’-CAG CAC TCT TTG AGG TGA AAA AC-3’, Reverse primer: 5’-ACG ATC TGA GAA CCA TTC GCA-3’; *Gpat3*, Forward primer: 5’-GGC CTT CGG ATT ATC CCT GG-3’, Reverse primer: 5’-CTT GGG GGC TCC TTT CTG AA-3’; *Plni2*, Forward primer: 5’-GAC CTT GTG TCC TCC GCT TAT-3’, Reverse primer: 5’-CAA CCG CAA TTT GTG GCT C-3’; *Plin3*, Forward primer: 5’-ATG TCT AGC AAT GGT ACA GAT GC-3’, Reverse primer: 5’-CGT GGA ACT GAT AAG AGG CAG G-3’; *Atp6v1b2*, Forward primer: 5’-ATG CGG GGA ATC GTG AAC G-3’, Reverse primer: 5’-AGG CTG GGA TAG GTA GTT CCG-3’; *Atp6ap2*, Forward primer: 5’-CTG GTG GCG GGT GCT TTA G-3’, Reverse primer: 5’-GCT ACG TCT GGG ATT CGA TCT −3’; *Dgat1*, Forward primer: 5’-TCC GTC CAG GGT GGT AGT G-3’, Reverse primer: 5’-TGA ACA AAG AAT CTT GCA GAC GA-3’; *Dgat2*, Forward primer: 5’-GCG CTA CTT CCG AGA CTA CTT-3’, Reverse primer: 5’-GGG CCT TAT GCC AGG AAA CT-3’; *mLpin1*, Forward primer: 5’-CAT GCT TCG GAA AGT CCT TCA-3’, Reverse primer: 5’-GGT TAT TCT TTG GCG TCA ACC T-3’; *Pnpla2*, Forward primer: 5’-GGA TGG CGG CAT TTC AGA CA-3’, Reverse primer: 5’-CAA AGG GTT GGG TTG GTT CAG-3’; *Lipe*, Forward primer: 5’-CCA GCC TGA GGG CTT ACT G-3’, Reverse primer: 5’-CTC CAT TGA CTG TGA CAT CTC G-3’; *Acsl1*, Forward primer: 5’-TGC CAG AGC TGA TTG ACA TTC-3’, Reverse primer: 5’-GGC ATA CCA GAA GGT GGT GAG-3’; *Acsl3,* Forward primer: 5’-AAC CAC GTA TCT TCA ACA CCA TC-3’, Reverse primer: 5’-AGT CCG GTT TGG AAC TGA CAG-3’; *Il-6*, Forward primer: 5’-AGT TGC CTT CTT GGG ACT GA-3’, Reverse primer: 5’-TCC ACG ATT TCC CAG AGA AC-3’; *β-actin*, Forward primer: 5’-AGC CAT GTA CGT AGC CAT CC-3’, Reverse primer: 5’-TCC CTC TCA GCT GTG GTG GTG AA-3’.

### Immunocytochemistry (ICC)

ICC was conducted by placing sterile glass coverslips in 12-well tissue culture plates and seeding approximately 2×10⁵ cells per well during their logarithmic growth phase. Upon reaching 70-80% confluency, cells were processed by first removing the culture medium and performing three washes with ice-cold PBS. Cellular fixation was achieved using 4% PFA in PBS at room temperature for 30 minutes, followed by three additional PBS washes. Permeabilization was then performed with 0.1% Triton X-100 in PBS for 10 minutes at room temperature, after which cells were washed three times with PBS. To reduce nonspecific binding, samples were blocked with 5% bovine serum albumin (BSA) in PBS for 1-2 hours at room temperature. Primary antibody incubation was carried out overnight at 4°C using antibodies diluted 1:500 in blocking buffer (5% BSA in PBS). The following day, cells underwent three 5-minute washes with PBST, followed by incubation with fluorophore-conjugated secondary antibodies (1:1000 dilution in blocking buffer) for 1 hour at room temperature under light-protected conditions. After three final 5-minute PBST washes, coverslips were mounted on glass slides using 6 μL of DAPI and allowed to air-dry protected from light before visualization using a laser scanning confocal microscope.

Secondary antibodies: Goat anti-Rabbit Alexa Fluor™ 488 (Thermofisher, A-11008), Donkey anti-Rabbit Alexa Fluor™ 647 (Thermofisher, A-31573).

### DQ-BSA assay

Log-phase cells were seeded onto glass coverslips in 24-well plates and cultured until reaching 80% confluence. For DQ-BSA staining, cells were incubated with 10 μg/mL DQ-BSA (prepared in complete growth medium) for 6-12 hours. Post-incubation, cells were washed three times with PBS and fixed with 4% PFA in PBS at room temperature for 30 min. Fixed samples were then washed three times with PBS with gentle agitation on an orbital shaker. For nuclear counterstaining, coverslips were mounted using 6 μL of DAPI, and imaged using a Leica SP8 confocal microscope.

### Lysotracker staining

For Lysotracker staining experiments, cells were seeded into confocal imaging dishes 24 hours prior and cultured until reaching 80% confluence. Immediately before staining, a working solution was prepared by diluting the 1 mM Lysotracker Red (1:20,000, Thermo Fisher, L12492) stock solution to 50 nM in complete culture medium pre-equilibrated to 37°C, while simultaneously adding Hoechst (1:10,000, Beyotime, P0133) to a final concentration of 1 μg/mL. The existing culture medium was carefully aspirated and replaced with the dye-containing working solution, followed by incubation at 37°C in 5% CO₂ for exactly 30 minutes. Post-incubation, cells were gently washed three times with pre-warmed PBS for 5 minutes per wash. Live-cell imaging was immediately performed using a laser scanning confocal microscope system equipped with appropriate filter sets for both dyes.

### Microglial conditioned medium (MCM) preparation and treatment protocols

The MCM was collected and centrifuged at 1,000 × g for 3 minutes at 4°C to remove cellular debris. The clarified supernatant was then filtered through a 0.45 μm sterile (Millipore, HAWP04700) membrane and aliquoted for subsequent experiments. siRNA-treated MCM (siRNA MCM): After plating, *Grn*^-/-^ BV2 microglial cells were transfected with 20 nM siRNA. The conditioned medium was collected 48 hours post-transfection, followed by identical processing steps.

Drug-treated MCM (FSG67 MCM): *Grn*^-/-^ BV2 microglial cells were plated and treated with varying concentrations of FSG67 (specific concentrations to be defined). After 48-hour treatment, the medium was harvested and processed identically (centrifugation and sterile filtration) to the standard MCM protocol.

### Cell viability and cytotoxicity assay

For CCK-8 proliferation assays, cells were plated in 96-well flat-bottom tissue culture plates at an optimal density of 2,000-3,000 viable cells per well and allowed to adhere overnight in a humidified 37°C incubator with 5% CO₂. Following complete attachment, the culture medium was carefully aspirated and replaced with phenol red-free DMEM supplemented with 10% CCK-8 reagent (MeilunBio, MA028-6). The plates were then incubated under standard culture conditions (37°C, 5% CO₂, 95% humidity) for exactly 2 hours, after which the absorbance was measured at 450 nm using a multimode microplate reader.

### Reactive oxygen species fluorescent (ROS) detection

For ROS detection, cells were plated in 96-well plates at a density of 2,000-3,000 viable cells per well (counted by trypan blue exclusion) and maintained in a humidified 37°C incubator with 5% CO₂ until reaching optimal confluency (typically 70-80%). The culture medium was carefully aspirated and replaced with 100 μL per well of 2’,7’-dichlorodihydrofluorescein diacetate (DCFH-DA, Beyotime, S0033S) working solution (10-20 μM final concentration, freshly prepared in serum-free and phenol red-free medium). For larger culture dishes, volumes were adjusted proportionally based on growth surface area (e.g., 2 mL for 35 mm dishes). Following gentle rocking to ensure uniform probe distribution, cells were incubated with the fluorescent probe for exactly 30 minutes at 37°C under light-protected conditions. After incubation, the DCFH-DA solution was completely removed by aspiration, and cells were washed 2-3 times with pre-warmed PBS (pH 7.4, 5 minutes per wash with gentle shaking) to thoroughly eliminate residual extracellular probe. Fluorescence intensity (excitation 485 nm/emission 535 nm) was quantified using a calibrated multimode microplate reader (TECAN SPARK) within 30 minutes of washing.

### SiRNA transfection

Three siRNAs targeting murine *Gpat3* were purchased from Biotend (catalog number S101). The sequences of siRNAs are: *Gpat3* siRNA1 sense: 5’-GCA CCA UCC AUU ACC AUA ATT-3’, antisense: 5’-UUA UGG UAA UGG AUG GUG CTT-3’; *Gpat3* siRNA2 sense: 5’-CCU CCU CAC ACG AAC CAA UTT-3’, antisense: 5’-AUU GGU UCG UGU GAG GAG GTT-3’; *Gpat3* siRNA3 sense: 5’-GCU ACC UGC UUC GAA UCA UTT-3’, antisense: 5’-AUG AUU CGA AGC AGG UAG CTT-3’; Negative control siRNA sense: 5’-UUC UCC GAA CGU GUC ACG UTT-3’, antisense: 5’-ACG UGA CAC GUU CGG AGA ATT-3’. Cells were plated in 6-well tissue culture plates and maintained until reaching approximately 60% confluence, the optimal density for transfection. To prepare the transfection mixture, 8.5 μL of sterile transfection buffer was first transferred to a nuclease-free microcentrifuge tube, followed by the addition of 0.75 μL of 20 μM siRNA stock solution (prepared in DEPC-treated, RNase-free water) to deliver a final amount of 15 pmol siRNA per well, with gentle vortexing to ensure complete homogenization. The transfection complex was then formed by adding 1.5 μL of PLUS™ Transfection Reagent (Shanghai Biotend Biotechnology Co., Ltd, TR-104-1000) to the siRNA mixture, followed by immediate thorough mixing via pipetting. The resulting siRNA-PLUS™ complexes were carefully administered dropwise to the cultured cells, and the plate was gently agitated to facilitate even distribution across the monolayer. Following a 72-hour incubation period under standard culture conditions (37°C, 5% CO₂ in a humidified atmosphere), the transfected cells were collected for downstream applications.

### Immunofluorescence and BODIPY/LipidSpot staining

Twenty-four hours prior to fixation, cells were plated on sterile glass coverslips in 24-well culture plates. Upon reaching 80-90% confluency, cells were fixed and subjected to standard immunofluorescence staining procedures. For simultaneous lipid staining, coverslips were co-incubated with secondary antibodies and either BODIPY 493/503 (1:1000, Thermofisher, D3922) or LipidSpot™ 610 (1:1000, Biotium, 70069-T) fluorescent dyes. The staining procedure was performed for 60 minutes at room temperature on an orbital shaker with light protection. Post-staining, coverslips were mounted in DAPI-supplemented antifade medium (ProLong™ Diamond) and visualized using a high-resolution confocal laser scanning microscope (Nikon NSPARC).

### Immunohistology and analysis

Immediately after perfusion, one cerebral hemisphere was collected, fixed in 4% PFA at 4 °C for 24 h, and then cryoprotected in 30% sucrose at 4 °C for 12-16 h until fully infiltrated. After briefly drying the surface, tissues were embedded in OCT and coronally sectioned at 30 μm using a cryostat at −20 °C. Sections were stored at −20 °C in cryoprotectant solution until use. For immunofluorescence staining, tissue sections were equilibrated to room temperature for 30 min, mounted on positively charged slides, and blocked with a solution of 5% BSA and 0.3% Triton X-100 in PBS for 1 h at 25°C in a humidified chamber. Following blocking, tissue sections were incubated with primary antibodies (overnight at 4°C) and corresponding secondary antibodies (1–2 hours at room temperature), washed three times with PBS, and finally imaged using a high-resolution laser-scanning confocal microscope. Antibodies used in this experiment are: PGRN (R&D, AF2557, 1:1,000), LAMP1 (Abcam, ab208943, 1:1,000), GPAT3 (Proteintech, 20603-1-AP, 1:2,000), IBA1 (Wako, 019-19741, 1:2,000), GFAP (Cell Signaling Technology, 12389, 1:1,000).

### Stereotaxic AAV injection

Mice were anesthetized with a safe dose of avertin and maintained on a heating pad at 37°C during and after surgery. Stereotaxic coordinates for the Posterior thalamic nucleus were determined with reference to Paxinos and Franklin’s The Mouse Brain in Stereotaxic Coordinates, 5th edition (Po: AP −2.06, ML±1.25, DV −3.2 relative to bregma). AAV-MG1.2-mCx3cr1-eGFP and AAV-MG1.2-mCx3cr1-mGrn-eGFP (FUBIO, 1×10¹³ vg/mL) were injected slowly using an automated stereotaxic apparatus (RWD, 71001-S). The needle was retained for 10 minutes post-injection, the scalp was sutured and disinfected, and animals were monitored post-operatively until recovery. Mice were retained for subsequent experiments only if body weight and general health remained stable.

### Intraperitoneal administration of FSG67 in mice

FSG67 (MCE, Cat. No HY-112489) was administered via intraperitoneal (i.p.) injection in mice under strict aseptic conditions. The drug was equilibrated to room temperature and visually inspected for precipitation prior to administration. Sterile syringes were loaded with precise dosages after complete air bubble removal. Mice were manually restrained in a supine position to expose the abdomen. The injection site (1-2 mm lateral to the lower abdominal midline) was disinfected with 70% alcohol or povidone-iodine swabs and air-dried. The needle was inserted at a 45° angle, advanced 3-5 mm subcutaneously before peritoneal cavity entry. After negative aspiration confirmation, FSG67 was injected slowly, followed by a 5-second pressure application with sterile gauze at the injection site.

Administration records documented injection timepoints, dosages, and treatment sequences. Mice underwent immediate post-injection monitoring followed by daily assessments of behavioral parameters (locomotor activity, neurological status) and physiological metrics (food/water consumption, fecal consistency). The treatment protocol comprised 15 administrations of FSG67 (2 mg/kg/dose) at 48-hour intervals, with all subsequent injections performed identically to the initial procedure.

### Novel object recognition test

Two identical cylindrical objects were placed at equivalent positions near the arena edges in a 40 × 40 cm box. Mice were placed facing away from the objects at equal distance and allowed to explore for 10 min (familiarization). One hour later, mice were reintroduced into the arena where one familiar object had been replaced by a novel object different in color, material and height. Interaction with novel versus familiar objects was recorded by SuperMaze to assess working memory.

### Light-dark transition test

The apparatus comprised a dark compartment (18 × 27 × 27 cm; enclosed; low illumination) and a light compartment (27 × 27 × 27 cm; open; higher illumination) connected by an opening. Animals were placed in the center of the light compartment facing away from the opening and allowed to explore for 10 min. Behavioral activity was recorded and analyzed with SuperMaze. An “entry” was scored when all four limbs entered a compartment.

### Open field test

Mice were placed in the lower-left corner of a 40×40 cm white arena and allowed to explore freely for 10 minutes. Behavioral experiments for Fig. 5h-i were recorded and analyzed using SuperMaze software, while those for Fig. 5l-m were recorded and analyzed using AnyMaze software. The central zone was defined as a 20 × 20 cm square; measures of total locomotion and center exploration were used to assess locomotor ability and anxiety-like behavior.

### RNA-seq analysis

For RNA sequencing analysis comparing WT BV2 and *Grn*^-/-^ BV2 cells (KO2 and KO8 clones) (three biological replicates per group), cells were cultured to a density of ≥1×10⁶ cells per extraction (optimal range: 3×10⁶–1×10⁷ cells), microscopically assessed for viability, and washed twice with RNase-free 1× PBS (room temperature, 1 min/wash). Cell lysis was performed using ice-cold RIPA buffer, and lysates were aliquoted into cryogenic tubes (certified for −192°C) for storage at −80°C or dry ice shipment. Total RNA was extracted using TRIzol^TM^ reagent (ThermoFisher,15596018CN), with integrity verified by Agilent Bioanalyzer (RIN ≥7). Poly(A)^+^ RNA was enriched via oligo(dT) magnetic beads, fragmented to 200–300 bp, and processed for strand-specific library preparation using dUTP marking. Libraries were quantified by qPCR, pooled at equimolar concentrations based on Bioanalyzer profiles, and sequenced on an Illumina NovaSeq platform (150 bp paired-end). Bioinformatics analysis included: (1) raw read QC (FastQC v0.11.9),(2) alignment to the mm10 genome (STAR v2.7.9a), (3) transcript quantification (featureCounts v2.0.3), (4) differential expression analysis (DESeq2 v1.34.0; thresholds: *p* <0.05), (5) functional annotation (clusterProfiler v4.2.2 using GO/KEGG databases).

### Lipidomic analysis

WT BV_2_ and *Grn*^-/-^ BV2 cells (three biological replicates per group) were cultured to 80–90% confluence, assessed for viability by phase-contrast microscopy, and washed twice with ice-cold RNase-free 1× PBS (1 min per wash with gentle orbital shaking at 100 rpm) while maintained on ice. Cells were detached using a sterile cell scraper in chilled PBS, transferred to pre-cooled 15 mL conical tubes by tilting culture plates at a 45° angle, and pelleted by centrifugation (4°C, 300–500 g for 5 min). Cell pellets were immediately flash-frozen in liquid nitrogen for 60 min using cryovials rated for −192°C, then stored at −80°C or shipped on dry ice with temperature monitoring. For lipidomic profiling, LC-MS/GC-MS raw data (raw/d files) were converted to mzXML format using ProteoWizard’s MSConvert (v3.0.22123) with centroiding enabled. Data processing involved: (1) XCMS Online (v3.15.2) for peak detection (matchedFilter algorithm, FWHM=10) and retention time alignment (obiwarp method), (2) CAMERA (v1.54.0) for adduct/isotope annotation (mass tolerance=5 ppm), (3) metaX (v1.4.20) for MS1/MS2 data integration, matching against HMDB (2023.1) and METLIN (2022.12) databases with a 10-ppm mass error. Metabolite identification combined KEGG (Release 107.1) pathway annotations and Reactome (v82) network topology. Differential metabolites were identified through multivariate statistics (PLS-DA with VIP>1) and univariate testing (Welch’s t-test, p<0.05, FDR-corrected), with |log2FC|>2 thresholds, visualized using ggplot2-based volcano plots and pheatmap-generated heatmaps (z-score normalized). Metabolic pathway impact analysis was performed in MetaboAnalyst 5.0 (globaltest algorithm) with KEGG Mapper integration for pathway visualization.

### Statistical analysis

All experimental data are expressed as mean ± standard error of the mean (SEM) derived from at least three independent biological replicates. Statistical comparisons were performed using GraphPad Prism 10.0, employing two-tailed Student’s t-tests for pairwise comparisons and one-way ANOVA with Tukey’s post hoc test for multi-group analyses, with statistical significance defined as *p* < 0.05.

## Acknowledgments

1. C. Zhu is sponsored by Research Startup Funds of Fudan University, National Natural Science Foundation of China (No. 82271476 and No. 82071436), Shanghai Pujiang Program (No. 20PJ1401100), and The Program for Oriental Scholars of Shanghai Universities (Distinguished Professor) (No. TP2022050). B. Li is supported by National Natural Science Foundation of China (No. 82101502) and China Postdoctoral Science Foundation (No. 2021M690036). This work is also supported by the Breakthrough Plan of the Ministry of Education, China (JYB2025XDXM604), and the Innovative research team of high-level local universities in Shanghai. The funders had no role in study design, data collection and analysis, decision to publish, or preparation of the manuscript.

## Data Availability Statement

Raw bulk RNA-seq data has been deposited in the NCBI Sequence Read Archive under BioProject accession PRJNA1401836. All data reported in this paper will be shared by the corresponding author upon request. Any additional information required to reanalyze the data reported in this paper is available from the corresponding author upon request. Correspondence and requests for materials should be addressed to C.Z

## Author contributions

Conceptualization, C.Z., B.L.; methodology, C.Z., B.L., J.L., Y.S., H.G.; investigation, J.L., Y.S., H.G., Y.L. T.Y., W.L., Y.W., Y.G.; writing-original draft, J.L., Y.S., H.G.; writing-review and editing, C.Z., B.L., J.L., Y.S., H.G.; funding acquisition, C.Z., B.L.; supervision, C.Z., B.L.; resources, C.Z., B.L.

## Supplementary Information is available for this paper

## Supplemental information

**Figure S1.**
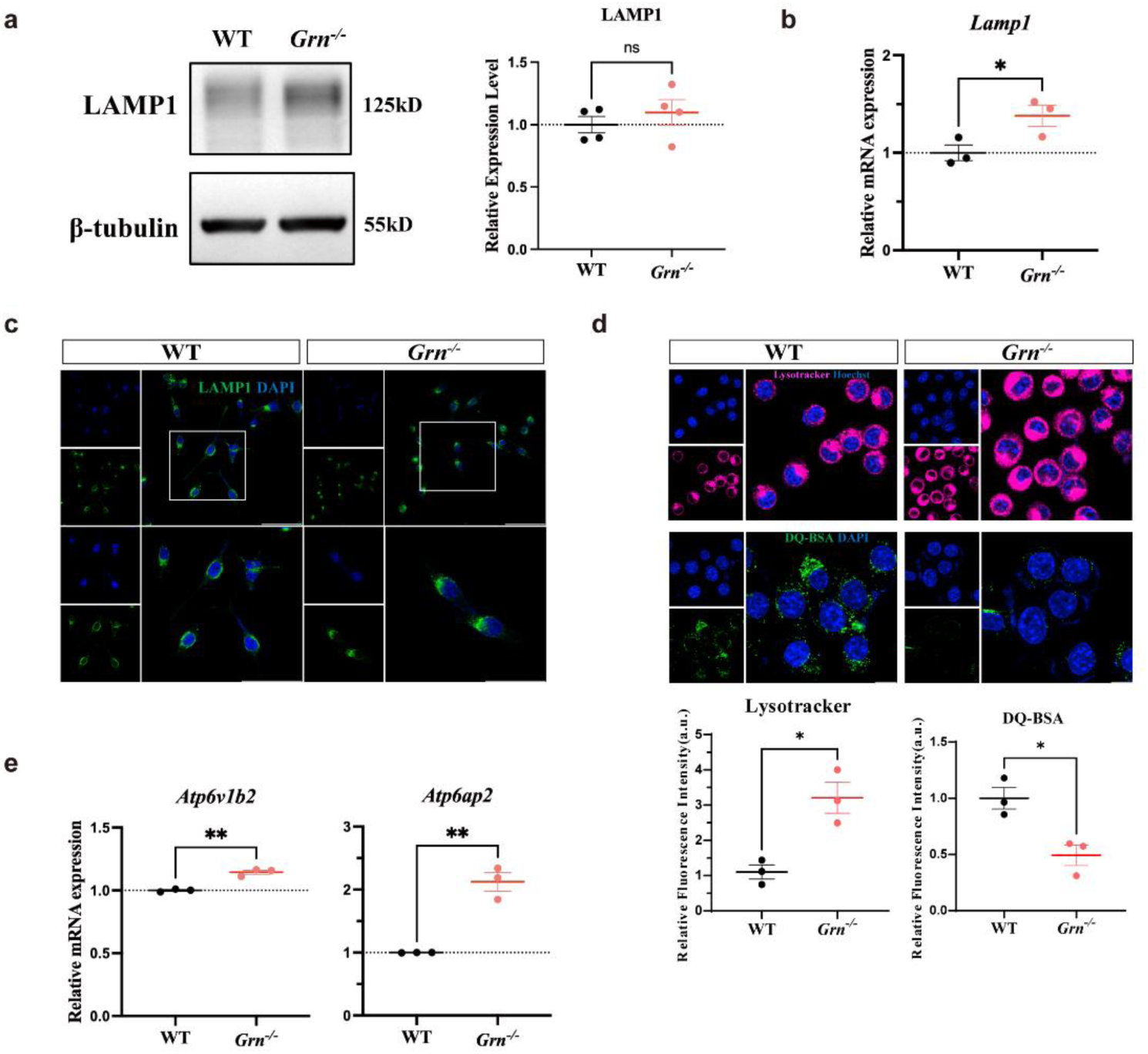
Ablation of *Grn* in microglia results in lysosomal dysfunction. a. Western blot analysis of LAMP1 and statistical results of gray value quantification. in *Grn^-/-^* BV2 microglial cells (n=4). b. Analysis of *Lamp1* mRNA levels in *Grn^-/-^* BV2 microglial cells (n=3). c. Representative LAMP1 fluorescence of *Grn^-/-^* BV2 microglial cells. Scale bar, 20 μm. d. Representative Lysotracker fluorescence and DQ-BSA staining of *Grn^-/-^*BV2 microglial cells and fluorescence quantification of Lysotracker signals (n = 3) and DQ-BSA signals (n = 3). Scale bar, 20 μm. e. Analysis of *Atp6v1b2* and *Atp6ap2* mRNA levels in *Grn^-/-^* BV2 microglial cells (n=3).

**Figure S2.**
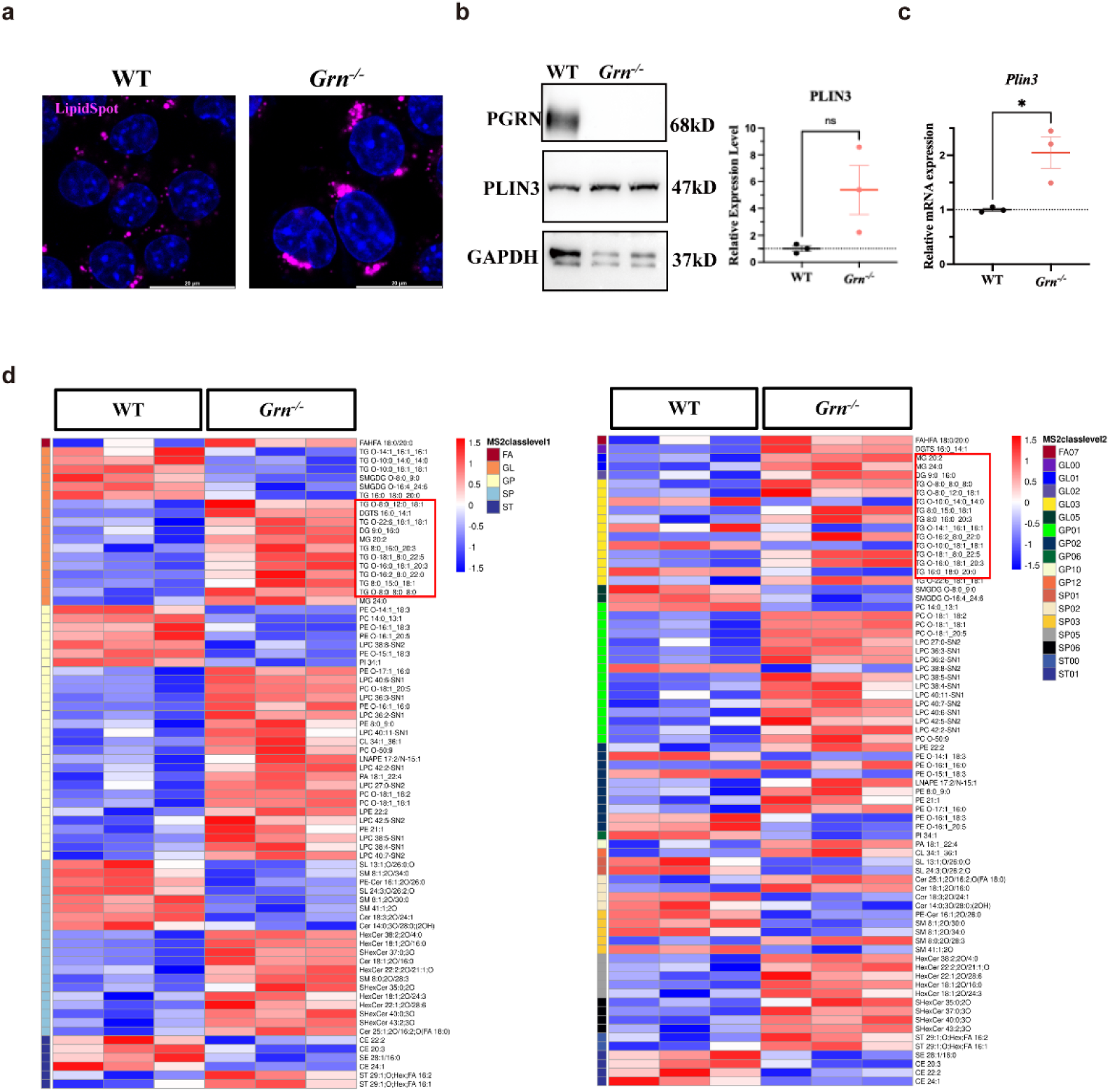
Lipid droplets accumulate in *Grn^-/-^* microglia stained by LipidSpot and lipidomic analysis of *Grn^-/-^* microglia. a. Representative LipidSpot staining of *Grn^-/-^* BV2 microglial cells. Scale bar, 20 μm. b. Western blot analysis of PLIN3 and statistical results of gray value quantification in *Grn^-/-^* BV2 microglial cells (n=3). c. Analysis of *Plin3* mRNA levels in *Grn^-/-^* BV2 microglial cells (n=3). d. Analysis of *Grn^-/-^* BV2 microglial cells lipidomic results. Clustering heat map analysis of differential lipids.

**Figure S3.**
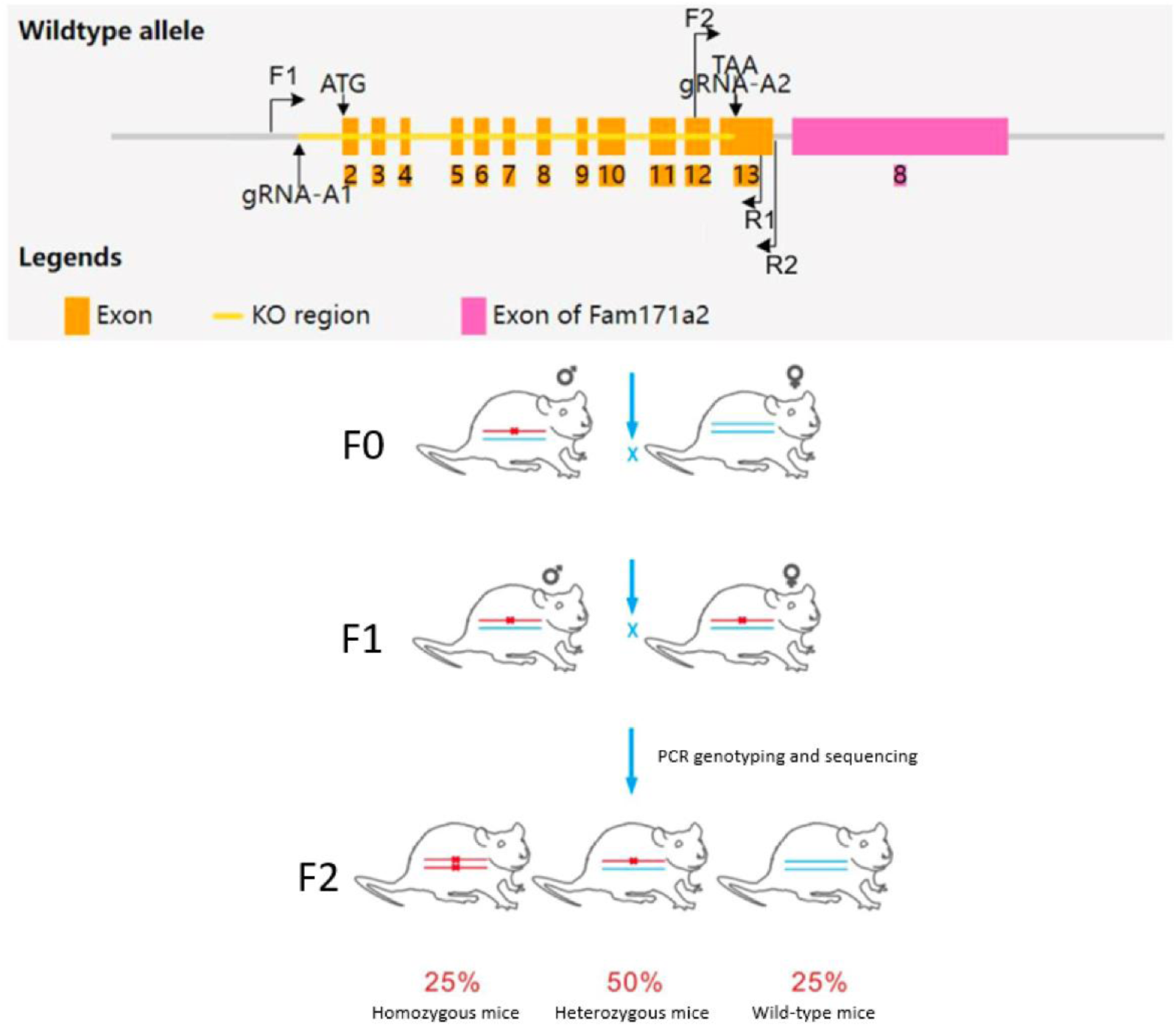
Schematic diagram of CRISPR-Cas9 based genome editing and mouse breeding strategy to generate *Grn* knockout mice.

**Figure S4.**
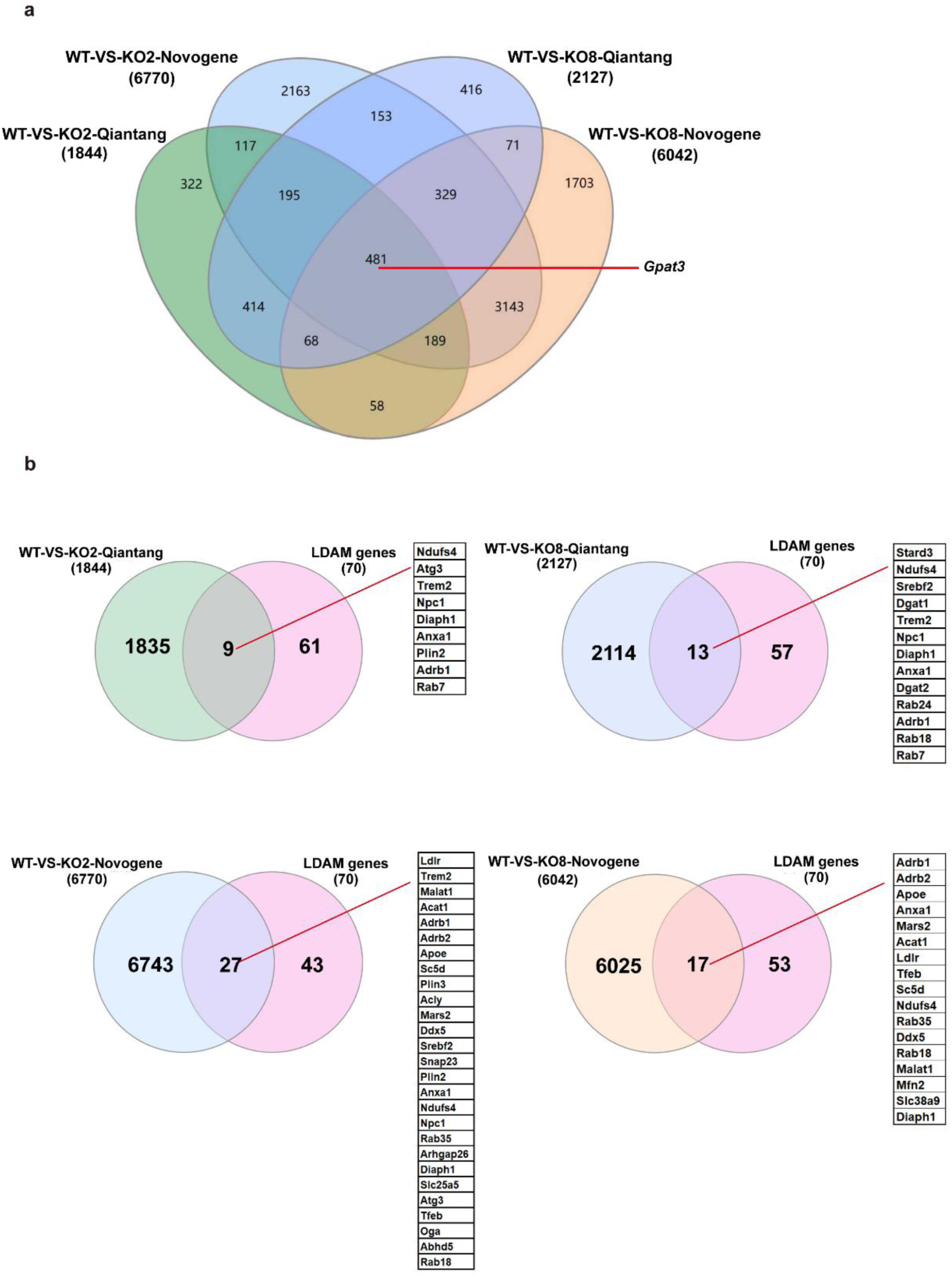
Venn diagram analysis of differentially expressed genes (DEGs) and their overlap with gene signatures related to lipid droplet (LD) biology. a. Venn diagram showing the overlapping of DEGs in four sets of transcriptomic comparison: WT vs. KO2 (Qiantang platform, 1844 DEGs), WT vs. KO2 (Novogene platform, 6770 DEGs), WT vs. KO8 (Qiantang platform, 2127 DEGs), and WT vs. KO8 (Novogene platform, 6042 DEGs). The unique and common DEGs among these comparisons are displayed. *Gpat3* is found in the intersection of the four groups. b. Four paired Venn diagrams illustrating the overlap of each individual dataset from Fig. S4a with the previously established LD signature genes. The left circle corresponds to the DEG set from Fig. S4a, and the right circle corresponds to the list of LD-related marker genes. The overlapping sections highlight shared genes between the DEGs in *Grn*^-/-^ BV2 microglial cells and the established LD-related genes.

**Figure S5.**
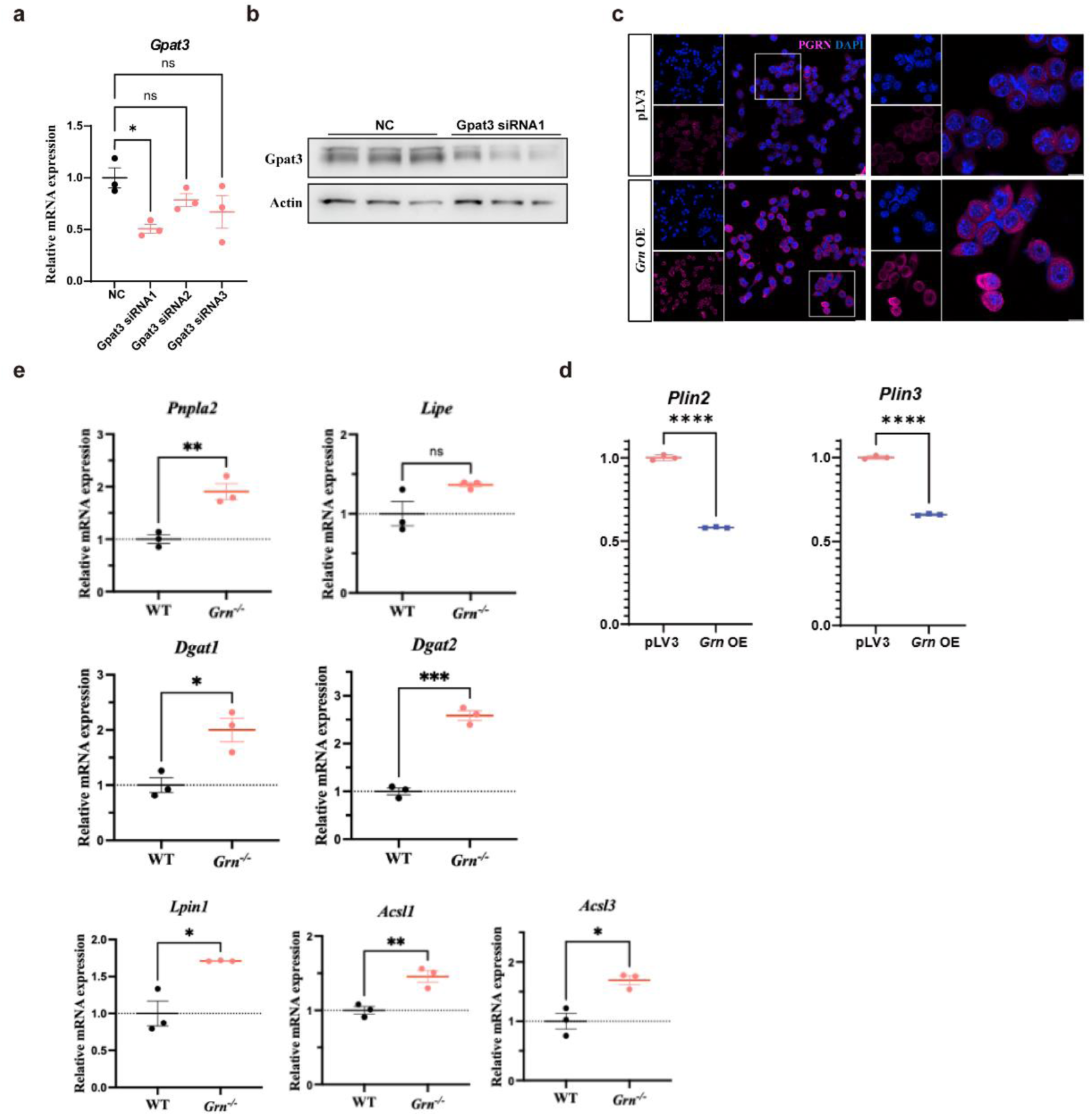
PGRN rescues lipid accumulation-induced neurotoxicity. a. Analysis of *Gpat3* mRNA levels in *Grn^-/-^* BV2 microglial cells and *Grn*^-/-^ BV2 cells treated with *Gpat3* siRNA1/2/3 (n=3). b. Western blot analysis of GPAT3 in control *Grn^-/-^* BV2 cells and *Gpat3* siRNA1-treated *Grn*^-/-^ BV2 cells (n=3). c. Representative PGRN fluorescence a of *Grn^-/-^* BV2 microglial cells and *Grn^-/-^ Grn* OE BV2 cells. Scale bar, 20 μm. d. Analysis of *Plin2* and *Plin3* mRNA levels in *Grn^-/-^* BV2 microglial cells and *Grn^-/-^Grn* OE BV2 cells (n=3). e. Analysis of enzymes for triglyceride degradation mRNA levels in *Grn^-/-^* BV2 microglial cells and *Grn^-/-^ Grn* OE BV2 cells (n=3).

**Figure S6.**
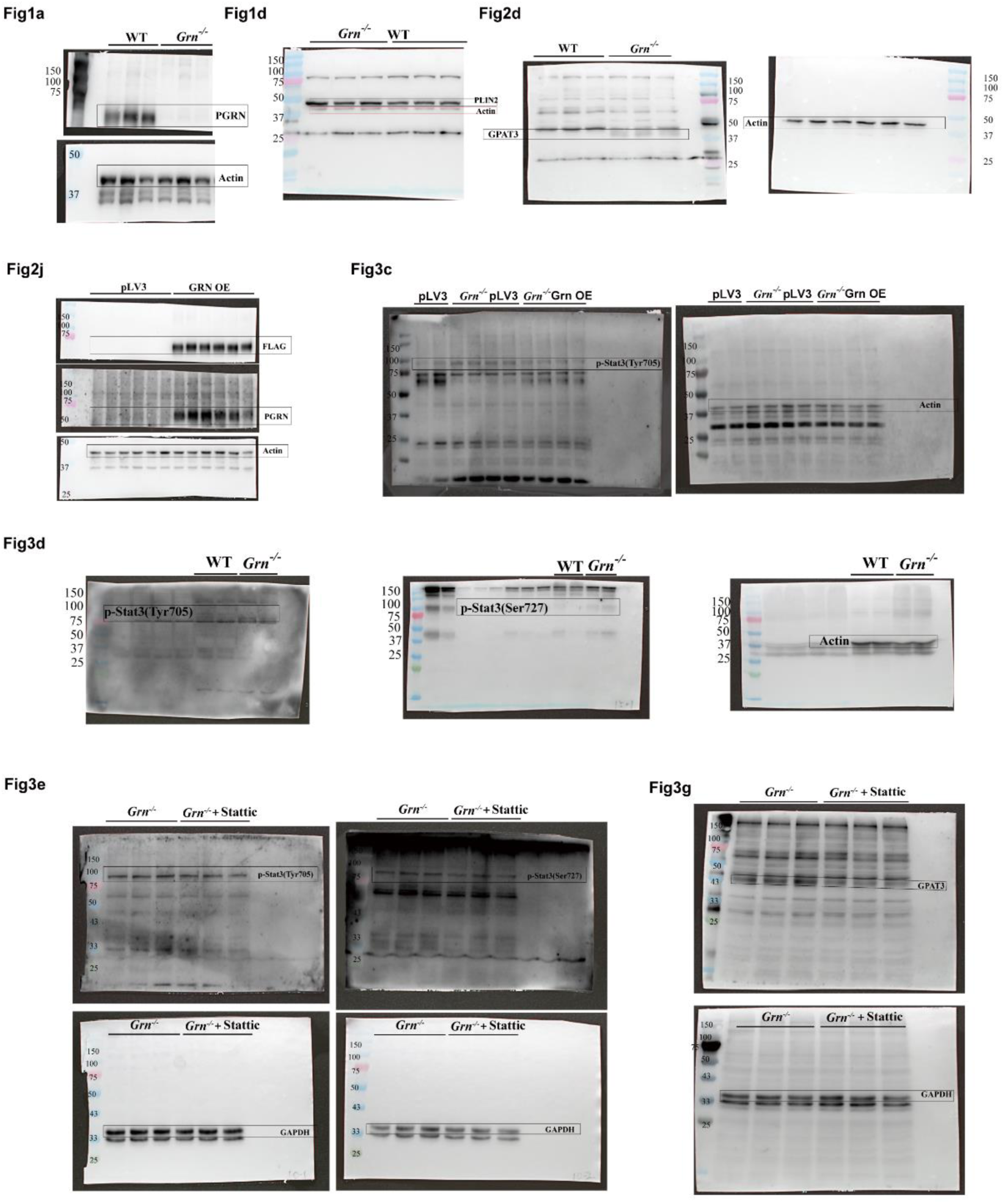
Uncropped scans of Western blots related to indicated figures.

**Table S1: Differentially expressed genes (DEGs) in *Grn* KO2 and KO8 cell lines analyzed by RNA-seq performed by Qiantang Biotech or Novogene.**

